# Targeting DNA topoisomerases or checkpoint kinases results in an overload of chaperone systems, triggering aggregation of a metastable subproteome

**DOI:** 10.1101/2021.06.04.447142

**Authors:** Wouter Huiting, Suzanne L. Dekker, Joris C. J. van der Lienden, Rafaella Mergener, Gabriel V. Furtado, Emma Gerrits, Maiara K. Musskopf, Mehrnoosh Oghbaie, Luciano H. Di Stefano, Maria A.W.H. van Waarde-Verhagen, Suzanne Couzijn, Lara Barazzuol, John LaCava, Harm H. Kampinga, Steven Bergink

## Abstract

A loss of the checkpoint kinase ATM leads to impairments in the DNA damage response, and in humans causes cerebellar neurodegeneration, and a high risk to cancer. A loss of ATM is also associated with increased protein aggregation. The relevance and characteristics of this aggregation are still incompletely understood. Moreover, it is unclear to what extent other genotoxic conditions can trigger protein aggregation as well. Here, we show that targeting ATM, but also ATR or DNA topoisomerases result in a similar, widespread aggregation of a metastable, disease-associated subfraction of the proteome. Aggregation-prone model substrates, including Huntingtin exon1 containing an expanded polyglutamine repeat, aggregate faster under these conditions. This increased aggregation results from an overload of chaperone systems, which lowers the cell-intrinsic threshold for proteins to aggregate. In line with this, we find that inhibition of the HSP70 chaperone system further exacerbates the increased protein aggregation. Moreover, we identify the molecular chaperone HSPB5 as a potent suppressor of it. Our findings reveal that various genotoxic conditions trigger widespread protein aggregation in a manner that is highly reminiscent of the aggregation occurring in situations of proteotoxic stress and in proteinopathies.

## INTRODUCTION

The PI3K-like serine/threonine checkpoint kinase ataxia telangiectasia mutated (ATM) functions as a central regulator of the DNA damage response (DDR), and is recruited early to DNA double-strand breaks (DSBs) by the MRE11/RAD50/NBS1 (MRN) complex (Shiloh and Ziv 2013). Defects in ATM give rise to ataxia-telangiectasia (A-T), a multisystem disorder that is characterized by a predisposition to cancer and progressive neurodegeneration (McKinnon 2012).

Impaired function of ATM has also been linked to a disruption of protein homeostasis and increased protein aggregation (Corcoles-Saez et al. 2018; Lee et al. 2018; Liu et al. 2005). Protein homeostasis is normally maintained by protein quality control systems, including chaperones and proteolytic pathways (Hipp, Kasturi, and Hartl 2019; Labbadia and Morimoto 2015). Together, these systems guard the balance of the proteome by facilitating correct protein folding, providing conformational maintenance, and ensuring timely degradation. When the capacity of protein quality control systems becomes overwhelmed during (chronic) proteotoxic stress, the stability of the proteome can no longer be sufficiently guarded, causing proteins to succumb to aggregation more readily. Proteins that are expressed at a relatively high level compared to their intrinsic aggregation-propensity, a state referred to as ‘supersaturation’, have been shown to be particularly vulnerable in this respect (Ciryam et al. 2015). A loss of protein homeostasis and the accompanying widespread aggregation can have profound consequences, and is associated with a range of (degenerative) diseases, including neurodegeneration (Kampinga and Bergink 2016; Klaips, Jayaraj, and Hartl 2018; Ross and Poirier 2004).

The characteristics and relevance of the aggregation induced by a loss of ATM are still largely unclear. Loss of MRE11 has recently also been found to result in protein aggregation (Lee et al. 2021), and since MRE11 and ATM function in the same DDR pathway, this raises the question whether other genotoxic conditions can challenge protein homeostasis as well (Ainslie et al. 2021; Huiting and Bergink 2021).

Here, we report that not just impaired function of ATM, but also inhibition of the related checkpoint kinase Ataxia telangiectasia and Rad3 related (ATR), as well as chemical trapping of topoisomerases (TOPs) using chemotherapeutic TOP poisons leads to widespread protein aggregation. Through proteomic profiling we uncover that the increased protein aggregation induced by these genotoxic conditions overlaps strongly with the aggregation observed under conditions of (chronic) stress and in various neurodegenerative disorders, both in identity and in biochemical characteristics. In addition, we find that these conditions accelerate the aggregation of aggregation-prone model substrates, including the Huntington’s disease related polyglutamine exon-1 fragment. We show that the widespread protein aggregation is the result of an overload of protein quality control systems, which can’t be explained by any quantitative changes in the aggregating proteins or by genetic alterations in their coding regions. This overload forces a shift in the equilibrium of protein homeostasis, causing proteins that are normally kept soluble by chaperones to now aggregate. Which proteins succumb to aggregation depends on the ground state of protein homeostasis in that cell-type. Finally, we provide evidence that the protein aggregation induced by genotoxic stress conditions is amenable to modulation by chaperone systems: whereas inhibition of HSP70 exacerbates aggregation, aggregation can be rescued by increasing the levels of the small heat shock protein HSPB5 (*α*B-crystallin).

## RESULTS

### Protein aggregation is increased upon targeting ATM, ATR or DNA topoisomerases

Aggregated proteins are often resistant to solubilization by SDS, and they can therefore be isolated using a step-wise detergent fractionation and centrifugation method. We isolated 1% SDS-resistant proteins (from here on referred to as aggregated proteins), and quantified these by SDS-PAGE followed by in-gel protein staining. In line with previous findings (Lee et al. 2018), we find that knocking out ATM in both U2OS and HEK293 results in an increase in protein aggregation (Figure 1A-C, Figure 1 – figure supplement 1A, B). Transient chemical inhibition of ATM (48-72 hours prior to fractionation) resulted in an increase in aggregated proteins in HEK293T cells as well (Figure 1C, D, Figure 1 – figure supplement 1C).

**Figure 1.**
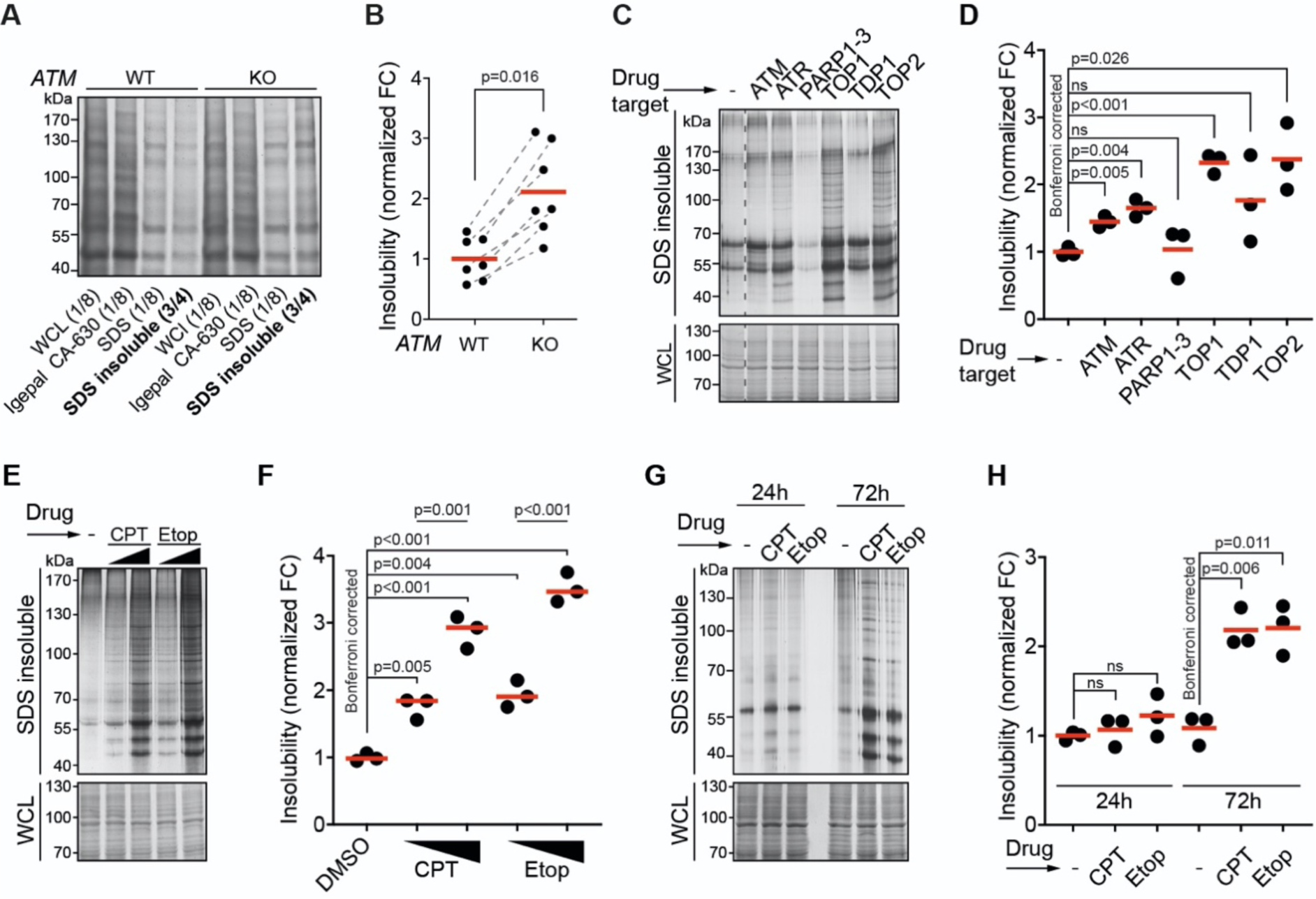
Protein aggregation is increased following a functional loss of ATM, ATR and upon topoisomerase poisoning. See also Figure 1 – figure supplement 1. (A) In-gel Coomassie staining of indicated fractions of cell extracts of WT and *ATM* KO U2OS cells. The relative amounts of each fraction loaded are indicated. (B) Quantification of A. Circles depict individual experiments; grey dotted lines depict matched pairs. Wilcoxon matched-pairs signed rank test, +/− standard deviation (C) Aggregated (silver stain) and whole cell lysate (WCL; Coomassie) fractions of HEK293T cells treated transiently with chemical agents targeting the indicating proteins (see Table 1 for drugs and doses used). (D) Quantification of C. Circles depict individual experiments. Two-tailed Student’s t-test with Bonferroni correction. (E) Protein fractions of HEK293T cells treated transiently with increasing amounts of camptothecin (CPT) (20-100 nM) or etoposide (Etop) (0.6-3 μM). (F) Quantification of E. Two-tailed Student’s t-test with Bonferroni correction. (G) Protein fractions of HEK293T cells treated transiently with CPT or Etop, targeting TOP1 or TOP2, respectively, 24 h or 72 h after treatment. (H) Quantification of G. Two-tailed Student’s t-test with Bonferroni correction.

**Table 1:** Antibodies used in this study

| **Primary** | **Application** | **Type** | **Supplier** | **Product ID** | **Concentration** |
| --- | --- | --- | --- | --- | --- |
| anti-GFP | IB | Mo,M | Takara Bio Clontech | 632380 | 1:5000 |
| anti-ATM | IB | Mo,M | Santa Cruz | sc-23921 | 1:200 |
| anti-HSPB5 | IB | Mo,M | Stressmarq | SMC-159 | 1:2000 |
| anti-GAPDH | IB | Mo,M | Fitzgerald | 10R-G109a | 1:10000 |
| anti-TUB | IB | Mo,M | Sigma | T5138 | 1:4000 |
| anti-HDAC1 | IB | Mo,M | DSHB | PCPR-HDAC1-2E12 | 0.5ug/ml |
| anti-MCM7 | IB | Mo,M | Santa Cruz | 47DC141 | 1:100 |
| anti-TUBA1A | IB | Mo,M | Sigma | T5168 | 1:2000 |
| anti-FUS | IF | Mo,M | Santa Cruz | sc-47711 | 1:200 |
| Anti-53BP1 | IF | Rb,P | Santa Cruz | sc-22760 | 1:150 |
| **Secondary** |  | **Type** | **Supplier** | **Product ID** | **Concentration** |
| anti-mouse | IB | Sh,P | GE Healthcare | NXA931 | 1:5000 |
| anti-rabbit | IB | Dk,P | GE Healthcare | NA934 | 1:5000 |
| anti-mouse,<br>Alexa fluor 488 | IF | Dk,P | Thermo Fisher | A-21202 | 1:500 |
| anti-rabbit,<br>Alexa fluor 488 | IF | Dk,P | Thermo Fisher | A-21206 | 1:500 |

Mo = mouse, Rb = rabbit, Dk = donkey, Sh = sheep; M = monoclonal, P = polyclonal; IB = immunoblotting, IF = immunofluorescence

Using the same experimental set-up, we examined the impact on aggregation of targeting various other DDR components. This revealed that chemical inhibition of the checkpoint signaling kinase ATR also enhanced protein aggregation. Inhibition of poly(ADP-ribose)polymerases 1-3 (PARP1-3), involved in single-strand break repair, or tyrosyl-DNA-phosphodiesterase 1 (TDP1), which repairs various 3’-blocking lesions including Topoisomerase 1 (TOP1) cleavage complexes, had no clear effect on protein aggregation (Figure 1C, D). This could be a result of functional redundancy, or in the case of TDP1, limited TOP1 trapping occurring under unstressed conditions in a timeframe of 72 hours. We therefore also directly targeted TOPs using the chemotherapeutic compounds camptothecin (CPT) and etoposide (Etop). The genotoxic impact of CPT and Etop is a well-documented consequence of their ability to trap (i.e. ‘poison’) respectively TOP1 and TOP2 cleavage complexes on the DNA, resulting in DNA damage (Pommier et al. 2010). Strikingly, we found that transient treatment with either compound caused a particularly strong increase in protein aggregation, which was dose-dependent (Figure 1C, D). Importantly, we observed no effect on aggregation within the first 24 hours after treatment with these compounds (Figure 1G, H). This reveals that the increased aggregation occurs only late, and argues that it does not stem from any immediate, unknown damaging effect of either CPT or Etop on mRNA or protein molecules. Together, these data indicate that the increased protein aggregation triggered by targeting ATM, ATR and topoisomerases is a late consequence of genotoxic stress.

### Camptothecin and ATM loss drive aggregation similarly, in a cell-type dependent manner

To investigate the nature of the proteins that become aggregated after genotoxic stress, we subjected the SDS-insoluble protein aggregate fractions and whole cell lysates (WCL) of CPT-treated HEK293T cells to label-free proteomics (Figure 2 – figure supplement 1A). We picked up a total of 983 aggregated proteins (Supplemental Table 1). Using a stringent cut-off (Benjamini-Hochberg corrected p<0.05; log_2_fold change>1; identified in >1 repeats of CPT-treated cells) (Figure 2A), we determined that 106 of these proteins aggregate significantly more after CPT-treatment (Figure 2A), compared to only 20 proteins that aggregate less. These 106 proteins aggregate highly consistent (Figure 2 – figure supplement 1B). Most of them were not identified as aggregating in untreated cells, implying that they are soluble under normal conditions (Figure 2 – figure supplement 1B). A GO-term analysis revealed that aggregating proteins are enriched for various cellular components, most notably mitochondria, myelin sheaths and ribonucleoprotein (RNP) complexes (Figure 2B). Many of them are RNA-binding proteins (Figure 2 – figure supplement 1C).

**Figure 2.**
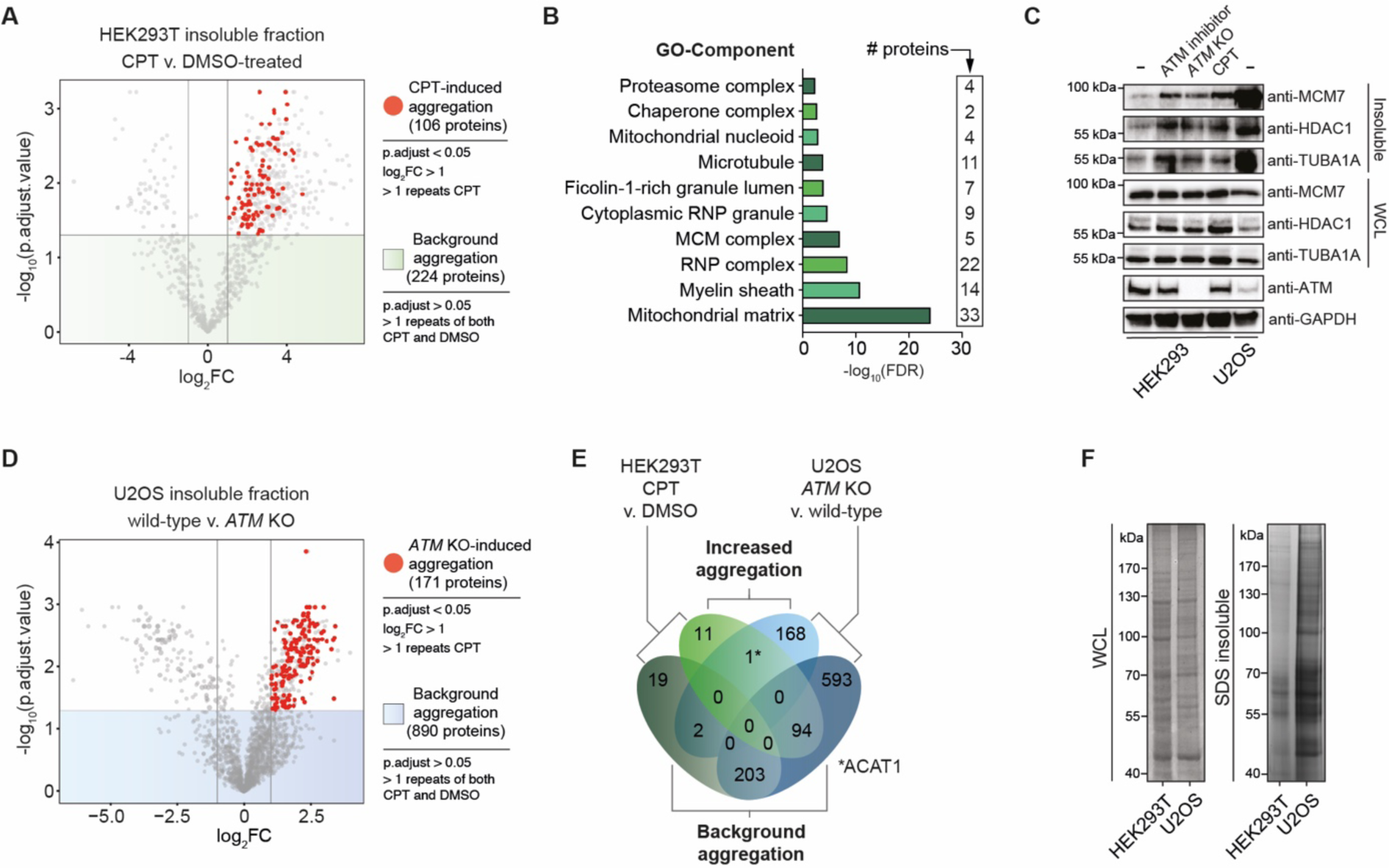
Topoisomerase poisoning and ATM loss have a highly similar, cell-type dependent impact on protein aggregation. See also Figure 2 – figure supplement 1. (A) Vulcanoplot of label-free quantification (LFQ) analysis of DMSO and CPT-treated HEK293T cells. n=4. (B) GO-term analysis (Component) of the increased aggregation in CPT-treated HEK293T cells. The top 10 terms with <2000 background genes are shown. (C) Western blot using the indicated antibodies on the aggregated and WCL fractions of drug-treated and *ATM* KO HEK239 cells, and wild-type U2OS cells. n=2. (D) Vulcanoplot of label-free quantification (LFQ) analysis of U2OS wild-type and *ATM* KO cells. n=4. (E) Venn diagram showing overlap between U2OS and HEK293T aggregation, for both background and increased aggregation. (F) Aggregated (silver stain) and whole cell lysate (WCL; Coomassie) fractions of untreated HEK293T and U2OS cells. n=2

Our initial silver stains suggested that the different drug treatments drive the aggregation of a similar set of proteins (Figure 1C). Indeed, a densitometry analysis revealed an almost identical staining pattern between them (Figure 2 – figure supplement 1D). Western blotting for MCM7, TUBA1A and HDAC1, three proteins that were identified in our MS analysis to aggregate more in CPT-treated HEK293T cells, confirmed that all three aggregated more in CPT-treated HEK293 cells. Importantly, MCM7, TUBA1A and HDAC1 also aggregated more than in unstressed conditions after treatment with ATM inhibitor, or when *ATM* was knocked out completely (Figure 2C). These findings indicate that these different genotoxic conditions have a similar impact on protein aggregation.

We therefore next investigated the aggregation caused by a loss of ATM in U2OS cells. Using the same MS pipeline, we identified a total of 1826 aggregated proteins across U2OS wild-type and *ATM* KO cells, almost twice as many as in HEK293T cells (Figure 2D, Supplemental Table 1). We found only 38 proteins that aggregated less in *ATM* KO cells, while 171 proteins aggregated significantly more. Of these 171 proteins, 91 were also found to aggregate more in *ATM*-depleted U2OS cells in a recent study by Lee *et al* (Lee et al. 2021). Similar to the CPT-induced aggregation in HEK293T cells, proteins that aggregate more in U2OS *ATM* KO cells appear to be largely soluble in wild-type cells, but now aggregate highly consistently (Figure 2 – figure supplement 1E).

However, despite the notion that different genotoxic conditions resulted in the aggregation of an overlapping set of proteins in HEK293 cells, at first glance, protein aggregation caused by a loss of *ATM* in U2OS cells seemed to be quite different. A GO-term analysis revealed limited overlap, with a loss of *ATM* in U2OS driving the aggregation of many microtubule and cytoskeleton (-related) components (Figure 2 – figure supplement 1F, G).

As protein aggregation can manifest vastly different in distinct cell-types (David et al. 2010; Freer et al. 2016), we examined which proteins already aggregate in the background of untreated HEK293T and U2OS cells (Benjamini-Hochberg corrected p>0.05, identified in >1 repeats of both case and control) (Figure 2A,D). This revealed a very strong overlap: 90% (203/225) of proteins that aggregated consistently – regardless of genotoxic stress – in HEK293T cells also consistently aggregated in U2OS cells (Figure 2E). Importantly, another 89% (94/106) of the proteins that aggregated more in CPT-treated HEK293T cells already aggregated in unstressed U2OS cells. This indicates that in U2OS cells a far bigger cluster of proteins ends up in aggregates, even under normal conditions. Indeed, silver staining revealed that in unstressed U2OS cells protein aggregation is substantially more prominent than in untreated HEK293T cells (Figure 2F). This is also reflected in MCM7, TUBA1A and HDAC1, all three of which aggregate strongly already in (untreated) wild-type U2OS cells (Figure 2C). These findings indicate that the lack of overlap between proteins that aggregate after CPT-treatment in HEK293T and proteins that aggregate in U2OS *ATM* KO cells is primarily a reflection of a different proteome and a different background aggregation in these two cell lines.

### Proteins that aggregate after genotoxic stress represent a metastable subproteome

Our data indicate that the genotoxic conditions of TOP1 poisoning and ATM loss have a comparable, but cell type-dependent impact on protein aggregation. In both HEK293T and U2OS cells, the protein aggregation does not appear to be limited to a specific location or function, but affects proteins throughout the proteome. This suggests that the aggregation is primarily driven by the physicochemical characteristics of the proteins involved.

A key determinant of aggregation is supersaturation. Protein supersaturation refers to proteins that are expressed at high levels relative to their intrinsic propensity to aggregate, which makes them vulnerable to aggregation. Supersaturation has been shown to underlie the widespread protein aggregation observed in age-related neurodegenerative disease, and in general ageing (Ciryam et al. 2015, 2019; Freer et al. 2019; Kundra et al. 2017; Noji et al. 2021). The relevance of supersaturation is underlined by the notion that evolutionary pressures appear to have shaped proteomes along its lines, so that at a global level, protein abundance is inversely correlated with aggregation propensity (Tartaglia et al. 2007). To determine the role of protein supersaturation in the aggregation observed in our experiments, we first defined a control group of proteins that were not identified as aggregating (NIA) in HEK293T cells, to serve as a benchmark. This group consisted of all proteins that were only identified in the HEK293T whole cell lysate, and not in the SDS-insoluble fraction. We next examined the intrinsic aggregation propensities of proteins, using the aggregation prediction tools TANGO (Fernandez-Escamilla et al. 2004) and CamSol (Sormanni and Vendruscolo 2019). Surprisingly, we found that aggregated proteins have in general a slightly lower (for background aggregation), or equal (for CPT-induced aggregation) intrinsic propensity to aggregate compared to NIA proteins (Figure 3A, Figure 3 – figure supplement 1A).

**Figure 3.**
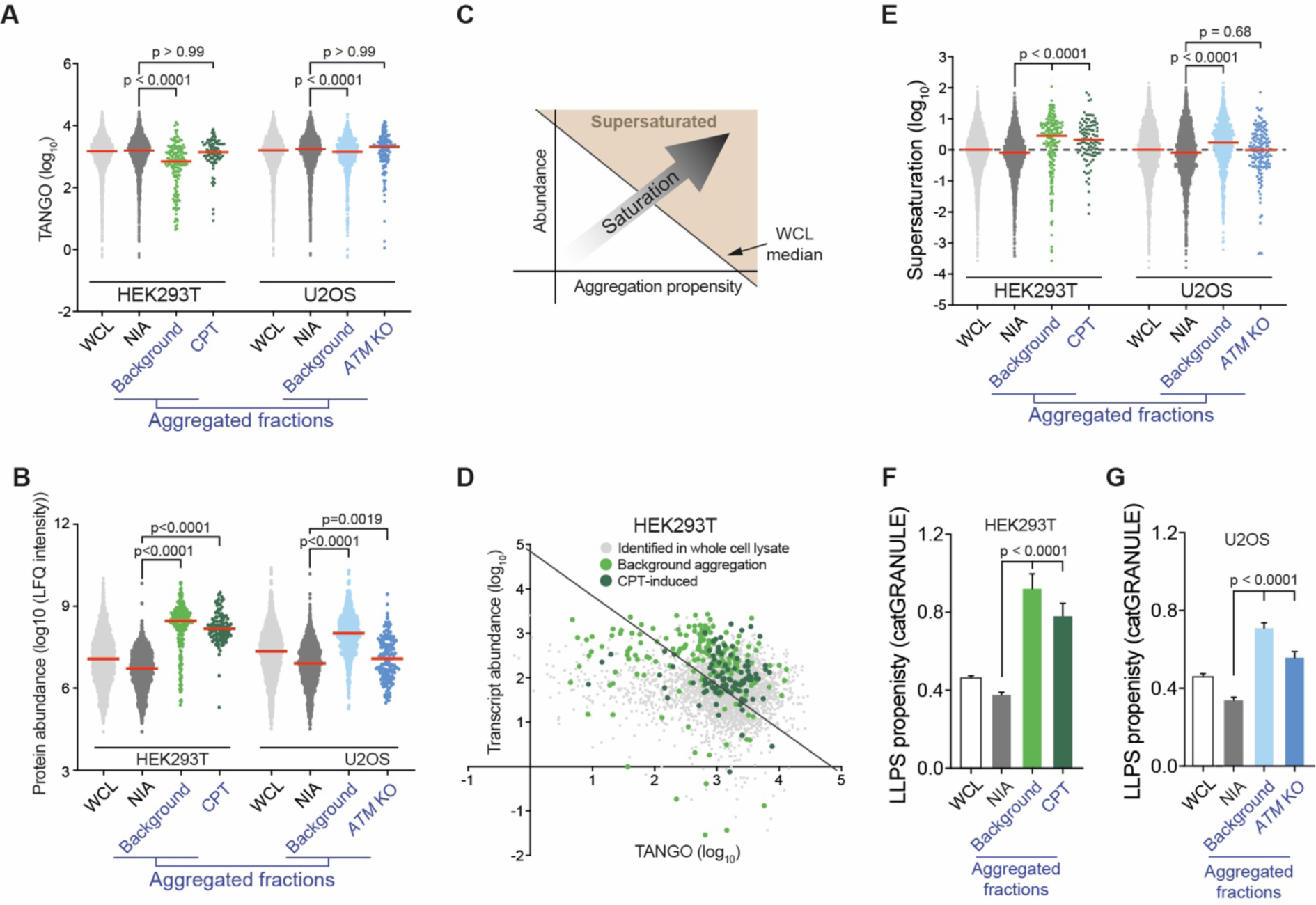
Proteins that aggregate after topoisomerase I poisoning are supersaturated and prone to engage in LLPS. See also Figure 3 – figure supplement 1. (A) TANGO scores of complete WCL, non-aggregated proteins (NIA) and aggregated fractions. (B) Protein abundance as measured by LFQ intensities. (C) Clarification of D. (D) Transcript abundances (as measured by RNAseq) plotted against TANGO scores, for the complete HEK293T MS analysis. All proteins above the diagonal (= HEK293T median saturation score, calculated using the HEK293T WCL dataset) are relatively supersaturated. (E) Supersaturation scores for the indicated protein fractions. (F,G) CatGRANULE scores for the indicated protein fractions. In all graphs, individual proteins and median values are shown. P-values are obtained by Kruskall-Wallis tests followed by Dunn’s correction for multiple comparisons.

However, even proteins with a low intrinsic propensity to aggregate can be supersaturated and be vulnerable to aggregation, when they are expressed at sufficiently high levels. Interestingly, our MS analysis revealed that proteins that aggregate in CPT-treated HEK293T cells are in general highly abundant compared to NIA proteins in (Figure 3B). Cross-referencing the aggregated proteins in our dataset against a cell-line specific NSAF reference proteome from Geiger *et al* (Geiger et al. 2012) (Figure 3 – figure supplement 1B) confirmed this.

To evaluate whether these proteins are indeed supersaturated, we used the method validated by Ciryam *et al,* which uses transcript abundance and aggregation propensity as predicted by TANGO to estimate supersaturation (Ciryam et al. 2013). After performing RNA sequencing on the same HEK293T cell samples that we used for our MS analysis (Figure 2 – figure supplement 1A, Supplemental Table 2), we confirmed that aggregating proteins are in general indeed more supersaturated than NIA proteins (Figure 3C,D). Cross-referencing our data against the composite human supersaturation database generated by Ciryam *et al* yielded a similar picture (Figure 3 – figure supplement 1E).

Although the relative supersaturation of aggregating proteins in HEK293T cells is intriguing, our data also indicates that most supersaturated proteins did not become SDS-insoluble, even after treatment with CPT (Figure 3D). Supersaturation only relates to overall protein concentration per cell, but within a cell, local protein concentrations can differ. A prime example of this is the partitioning of proteins in so-called biomolecular condensates through liquid-liquid phase separation (LLPS). LLPS can increase the local concentration of proteins, which has been shown to be important for a wide range of cellular processes (Lyon, Peeples, and Rosen 2021). However, it also comes with a risk of transitioning from a liquid to a solid, and even amyloid state. Indeed, a large amount of recent data have clearly demonstrated that proteins that engage in LLPS are overrepresented among proteins that aggregate in various proteinopathies (reviewed in Alberti and Hyman 2021). Using catGRANULE (Mitchell et al. 2013) (http://tartaglialab.com), we find that both background and CPT-induced aggregation are indeed made up of proteins that have a higher average LLPS-propensity than NIA proteins (Figure 3F). Both background and CPT-induced aggregation are also enriched for proteins that have a high propensity to engage in LLPS-relevant pi-pi interactions, as indicated by both a higher average PScore, and a larger percentage of proteins that have a PScore > 4 (i.e. above the threshold defined by Vernon et al., 2018) (Figure 3 – figure supplement 1F,G). Inversely, dividing NIA proteins into supersaturated and non-supersaturated subgroups reveals that they have a similarly low average LLPS-propensity (Figure 3 – figure supplement 1H,I). This points out that a high LLPS-propensity can discriminate supersaturated proteins that are prone to aggregate from supersaturated proteins that are not.

Upon examining the proteins that aggregate in U2OS cells, we found further support for this. Background aggregation in U2OS cells is also made up of supersaturated, LLPS-prone proteins. Despite this background aggregation being far more pronounced in U2OS cells than in HEK293T cells, many supersaturated proteins are not SDS-insoluble in U2OS cells, even in cells lacking *ATM* (Figure 3 – figure supplement 1C). In U2OS *ATM* KO cells, aggregating proteins are not more supersaturated per se than U2OS NIA proteins, nor do they have higher PScores (Figure 3A,B,E; Figure 3 – figure supplement 1A,C-E,J,K). However, they do have a higher general propensity to engage in LLPS as predicted by catGRANULE (Figure 3G). From this we conclude that both CPT-treatment and a loss of ATM further exacerbate the aggregation of LLPS-prone and supersaturated proteins, in a cell-type dependent manner.

### Genotoxic stress-induced protein aggregation is the result of a global lowering of the protein aggregation threshold

Our data shows that a substantial number of inherently similarly vulnerable proteins aggregate under the genotoxic conditions of CPT treatment or ATM loss. Their consistent aggregation across independent repeats argues against the possibility that this is caused by any genotoxic stress-induced DNA sequence alterations in their own coding regions, as these would occur more randomly throughout genome. Moreover, we find that the increased aggregation can also not be explained by any changes in abundance of the proteins involved, resulting for example from DNA damage-induced transcriptional dysregulation, as very limited overlap exists between proteins that aggregate and proteins with an altered expression upon CPT-treatment or ATM loss (Figure 4A).

**Figure 4.**
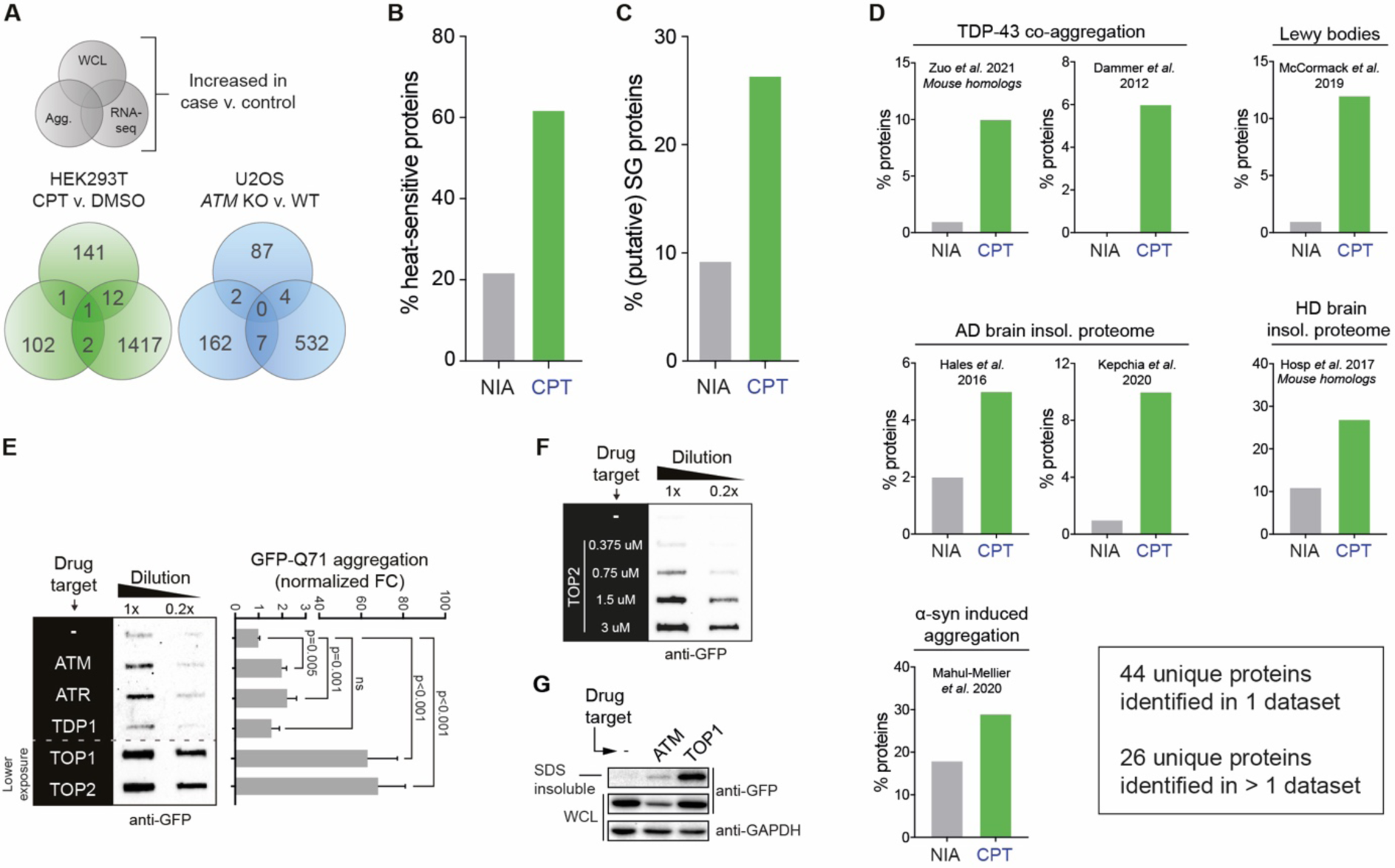
The cell-intrinsic aggregation threshold is lowered upon targeting ATM, ATR or DNA topoisomerases. See also Figure 4 – figure supplement 1. (A) Overlap between RNA sequencing analysis and LFQ MS analysis for WCL and aggregated protein fractions. Only significant changes are taken into account. (B) Relative occurrences of proteins that have been shown to aggregate upon heat-stress. See text for reference. (C) Relative occurrences of proteins that have been found to associated with stress granules. See text for reference. (D) Relative occurrences of aggregated proteins in various disease (model) datasets, obtained from the indicated studies. See also Figure 4 – figure supplement 1D. (E) Left panel: filter trap assay of HEK293 cells expressing inducible Q71-GFP that received the indicated treatment, probed with GFP antibody. n=3. Right panel: quantification, using Student’s two-tailed t-test followed by a Bonferroni correction for multiple comparisons. (F) Filter trap assay of HEK293 cells expressing inducible Q71-GFP that were treated with the indicated doses of Etop, probed with GFP antibody. n=2 (G) Western blot of WCL and aggregated proteins isolated from HEK293 cells expressing inducible luciferase-GFP, treated with ATM inhibitor or CPT, probed with the indicated antibodies. n=2

Instead, our data indicate that a long-term consequence of these genotoxic conditions is a global lowering of the aggregation threshold of proteins. As a result, more and more LLPS-prone, supersaturated proteins that are normally largely soluble now start to aggregate, with the most vulnerable proteins aggregating first. This aggregation threshold appears to be inherently lower in U2OS cells compared to HEK293T cells, causing a large population of metastable proteins to aggregate already under normal conditions. A loss of ATM in U2OS cells lowers the aggregation-threshold even further, causing a ‘second layer’ of LLPS-prone proteins that are not even supersaturated to aggregate also (Figure 4 – figure supplement 1A).

This lowering of the aggregation threshold is highly reminiscent of ‘classic’ protein aggregation resulting from (chronic) proteotoxic stresses (Weids et al. 2016), and has been referred to as a disturbed (Hipp et al. 2019) or shifted protein homeostasis (Ciryam et al. 2013). In line with this, we find that many of the proteins that aggregate after CPT treatment have also been reported to aggregate upon heat treatment of cells (Figure 4B) (Mymrikov et al. 2017). In addition, more than 30 of them have previously been found to associate with stress-granules (Figure 4C) (http://rnagranuledb.lunenfeld.ca), cellular condensates that have been found to function as nucleation sites for protein aggregation (Dobra et al. 2018; Mateju et al. 2017). A shift in protein homeostasis has also been suggested to be key to the build-up of protein aggregates during ageing (Ciryam et al. 2014) and to the initiation of protein aggregation in a range of chronic disorders (David et al. 2010; Hipp et al. 2019; Morley et al. 2002). Intriguingly, we find that proteins that aggregate after transient CPT-treatment are enriched for constituents of various disease-associated protein aggregates (Figure 4D). 68% (72/106) of them – or their mouse homologs – has already previously been identified in TDP-43 aggregates (Dammer et al. 2012; Zuo et al. 2021), Lewy bodies (McCormack et al. 2019), or *α*-synuclein induced aggregates (Mahul-Mellier et al. 2020), or found to aggregate in Huntington’s disease (Hosp et al. 2017) or Alzheimer’s disease brains (Hales et al. 2016; Kepchia et al. 2020) (Figure 4 – figure supplement 1B).

If genotoxic conditions indeed over time lead to a lowering of the aggregating threshold, this would predict that they can also result in an accelerated aggregation of aggregation-prone model substrates. For example, disease-associated expanded polyQ-proteins are inherently aggregation-prone, and they have been shown to aggregate faster in systems in which protein homeostasis is impaired (Gidalevitz et al. 2013; Gidalevitz, Kikis, and Morimoto 2010). We went back to HEK293 cells, and employed a line carrying a stably integrated, tetracycline-inducible GFP-tagged Huntingtin exon1 containing a 71 CAG-repeat (encoding Q71). Transient targeting of ATM, ATR and in particular topoisomerases, but not TDP1, 24-48 hours prior to the expression of polyQ (Figure 4 – figure supplement 1C) indeed accelerated polyQ aggregation in these cells (Figure 4E, Figure 4 – figure supplement 1D), closely mirroring the increased aggregation that we observed before (Figure 1C,D). The accelerated polyQ aggregation under these conditions is also dose-dependent (Figure 4F, Figure 4 – figure supplement 1E, F). PolyQ aggregation is normally proportional to the length of the CAG repeat, which is intrinsically unstable. Importantly, we find no evidence that the accelerated polyQ aggregation induced by these genotoxic conditions can be explained by an exacerbated repeat instability (Figure 4 – figure supplement 1G). Next, we also used the same tetracycline-inducible system and experimental set-up to investigate the aggregation of the protein folding model substrate luciferase-GFP (Figure 4 – figure supplement 1H). We find that transient targeting of either ATM or TOP1 results in an enrichment of luciferase-GFP in the aggregated fraction (Figure 4E).

### Genotoxic stress results in a rewiring of chaperone networks which is however insufficient to prevent client aggregation

We noted that CPT-treatment resulted in an increased aggregation of multiple (co)chaperones in HEK293T cells. We identified 17 aggregating (co)chaperones, 5 of which aggregate significantly more after treatment with CPT (Figure 5A). In U2OS cells many (co)chaperones are already aggregating in the background, but still a few aggregated significantly more in *ATM* KO cells (Figure 5B). These findings are interesting, as chaperone systems have the ability to modulate aggregation (Hartl, Bracher, and Hayer-Hartl 2011; Mogk, Bukau, and Kampinga 2018; Sinnige, Yu, and Morimoto 2020; Tam et al. 2006). HSP70s (HSPAs) are among the most ubiquitous chaperones, and they have been shown to play a key role in maintaining protein homeostasis in virtually all domains of life (Gupta and Singh 1994; Hunt and Morimoto 1985; Lindquist and Craig 1988). Upon cross-referencing the NIA and aggregating fractions against a recently generated client database of HSPA8 (HSC70; constitutively active form of HSP70) and HSPA1A (constitutively active and stress-inducible HSP70) (Ryu et al. 2020), we find that HSPA8 and HSPA1A clients are enriched among aggregating proteins (Figure 5C). We also mined the BioGRID human protein-protein interaction database using the complete KEGG dataset of (co)chaperones (168 entries). Although the transient and energetically weak nature of the interactions between many (co)chaperones and their clients (Clouser et al. 2019; Kampinga and Craig 2010; Mayer 2018) makes it likely that these interactions are underrepresented in the BioGRID database, it can provide additional insight into the presence of (putative) chaperone clients in the aggregating fractions (Victor et al. 2020). We find that all aggregating fractions, including U2OS *ATM* KO aggregation, are enriched for (co)chaperone interactors (Figure 5 – figure supplement 1A). Aggregating proteins have reported interactions with a broad range of chaperone families, most notably HSP70s and HSP90s (and known co-factors of these), and chaperonins (primarily TRiC/CCT subunits) (Figure 5D and Figure 5 – figure supplement 1B,C).

**Figure 5.**
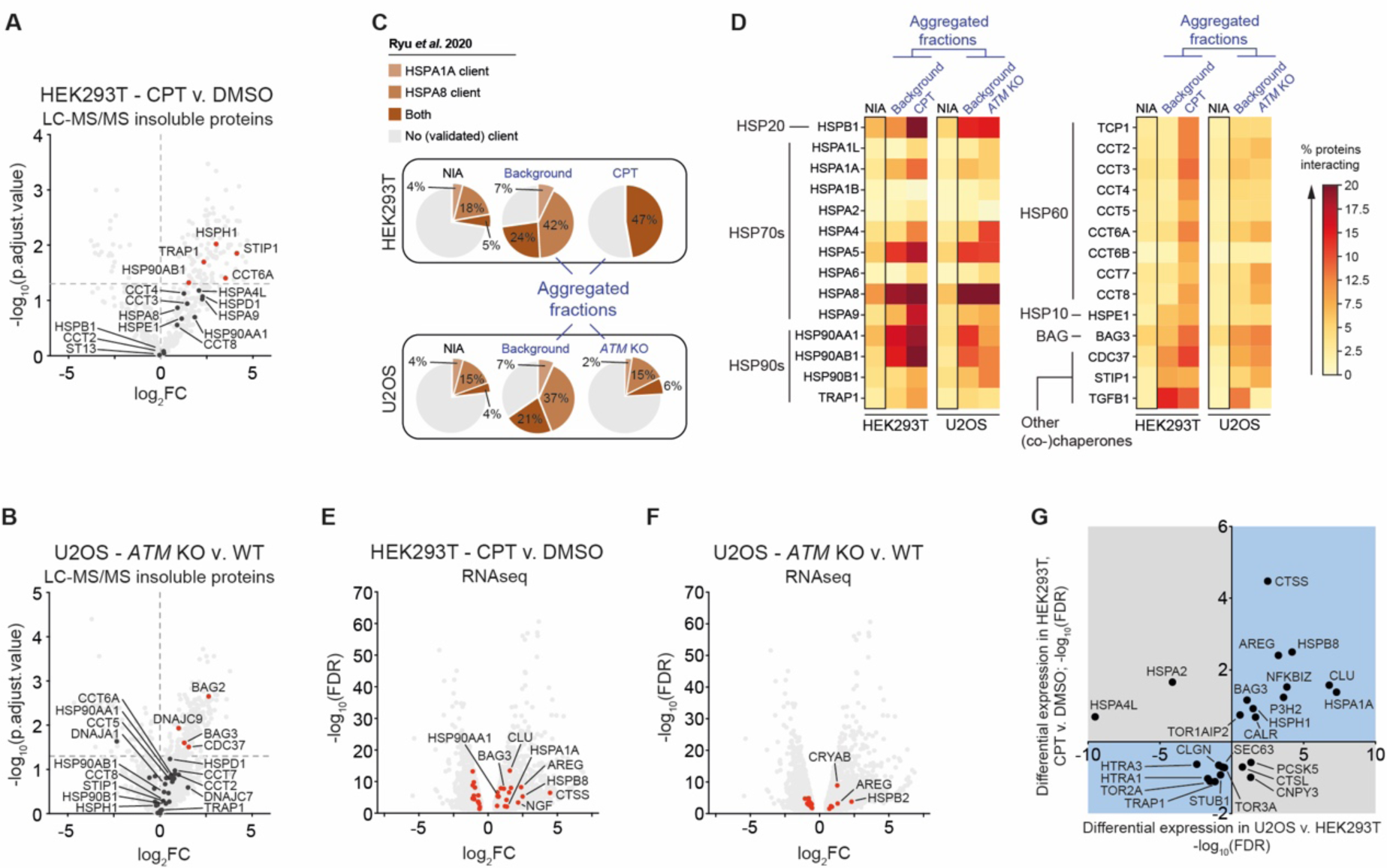
The lowered aggregation threshold caused by topoisomerase poisoning or a loss of ATM is accompanied by a rewiring and aggregation of known interacting (co)chaperones. See also Figure 5 – figure supplement 1. (A,B) Presence of (co)chaperones in the aggregated protein fractions in HEK293T and U2OS cells. (C) Pie charts showing the presence of HSPA1A and HSPA8 clients in aggregated protein fractions, compared to clients present in both NIA fractions. See text for reference; only clients identified in at least two out of three repeats were taken into account here. (D) BioGRID (co)chaperone interactions with the aggregated proteins identified in this study, per (co)chaperone. Darker colors represent a higher percentage of proteins with a reported binding to that (co)chaperone. See also Supplemental Figure 5B. (E) Differentially expressed (co)chaperones in CPT-treated HEK293T cells compared to DMSO-treated cells. (F) Differentially expressed (co)chaperones in U2OS *ATM* KO cells compared to wild-type cells. (G) Graphs showing (co)chaperones that are differentially expressed in both CPT-treated HEK293T cells compared to DMSO-treated HEK293T cells, and in untreated U2OS compared to untreated HEK293T cells.

Intriguingly, the (co)chaperones that we found to aggregate themselves are among the most frequent interactors. This suggests that they were sequestered by protein aggregates as they engaged their client proteins, in line with what has been reported for disease-associated aggregation (Hipp et al. 2019; Jana 2000; Kim et al. 2013; Mogk et al. 2018; Yu et al. 2019; Yue et al. 2021). Overall, we find that the relative levels of chaperone engagement of the different aggregating fractions largely reflect their respective supersaturation and LLPS-propensities.

When the capacity of chaperone systems is overloaded, this can trigger an up-regulation of chaperone levels. This plasticity of chaperone systems allows cells to adapt to varying circumstances and proteotoxic stress conditions (Klaips et al. 2018). In HEK293T cells, we find that treatment with CPT results in an overall upward shift of (co)chaperone expression levels, as measured in both our RNAseq dataset (16 up, 12 down) (Figure 5E), and our WCL MS analysis (15 up, 7 down) (Figure 5 – figure supplement 1D). Upregulated chaperones include HSPB1, DNAJA1, HSPA1A, HSPA5, HSPA8, HSP90AA1, and BAG3, all of which are among the most frequent interactors of aggregating proteins in CPT-treated cells. In U2OS cells, a loss of ATM appears to result in a more balanced rewiring of chaperone systems compared to wild-type cells (Figure 5F, Figure 5 – figure supplement 1E). Nevertheless, similar to HEK293T cells, many of the most frequent (co)chaperone interactors of the aggregating proteins in U2OS are found to aggregate themselves as well. These findings indicate that genotoxic stress induces a rewiring of chaperone systems, which is however insufficient to prevent the increased aggregation of metastable client proteins.

We reasoned that the difference in background aggregation between HEK293T and U2OS cells might also be reflected in different chaperone expression levels already under normal conditions. Indeed, a differential expression analysis between untreated HEK293T and untreated U2OS cells revealed a strong overall upward shift of (co)chaperone transcript levels in the latter (Figure 5 – figure supplement 1F). For example, we found that transcript levels of the small heat shock-like protein Clusterin (CLU) are >100-fold higher in wild-type U2OS compared to HEK293T cells, and that transcript levels of the stress-inducible HSPA1A are >150-fold higher. Interestingly, the differences in expression of chaperone systems in U2OS compared to HEK293T overlap with the changes occurring after CPT treatment in the latter. Out of the 24 (co)chaperones identified to be expressed differently in both, 19 are altered in the same direction (Figure 5G).

### Genotoxic stress-induced protein aggregation is amenable to modulation by chaperone systems

Our data suggest that the lowering of the aggregation threshold upon various genotoxic conditions is caused by an overload of chaperone systems, leading to a shift in protein homeostasis. We reasoned that targeting chaperone systems may then exacerbate aggregation. Indeed, mild HSP70 inhibition using the HSP70/HSC70 inhibitor VER-155008 after CPT-treatment increased CPT-induced protein aggregation even further, while having no clear impact on aggregation in control cells (Figure 6A). Similar results were obtained when we blotted the aggregated fractions for MCM7 and TUBA1A (Figure 6B).

**Figure 6.**
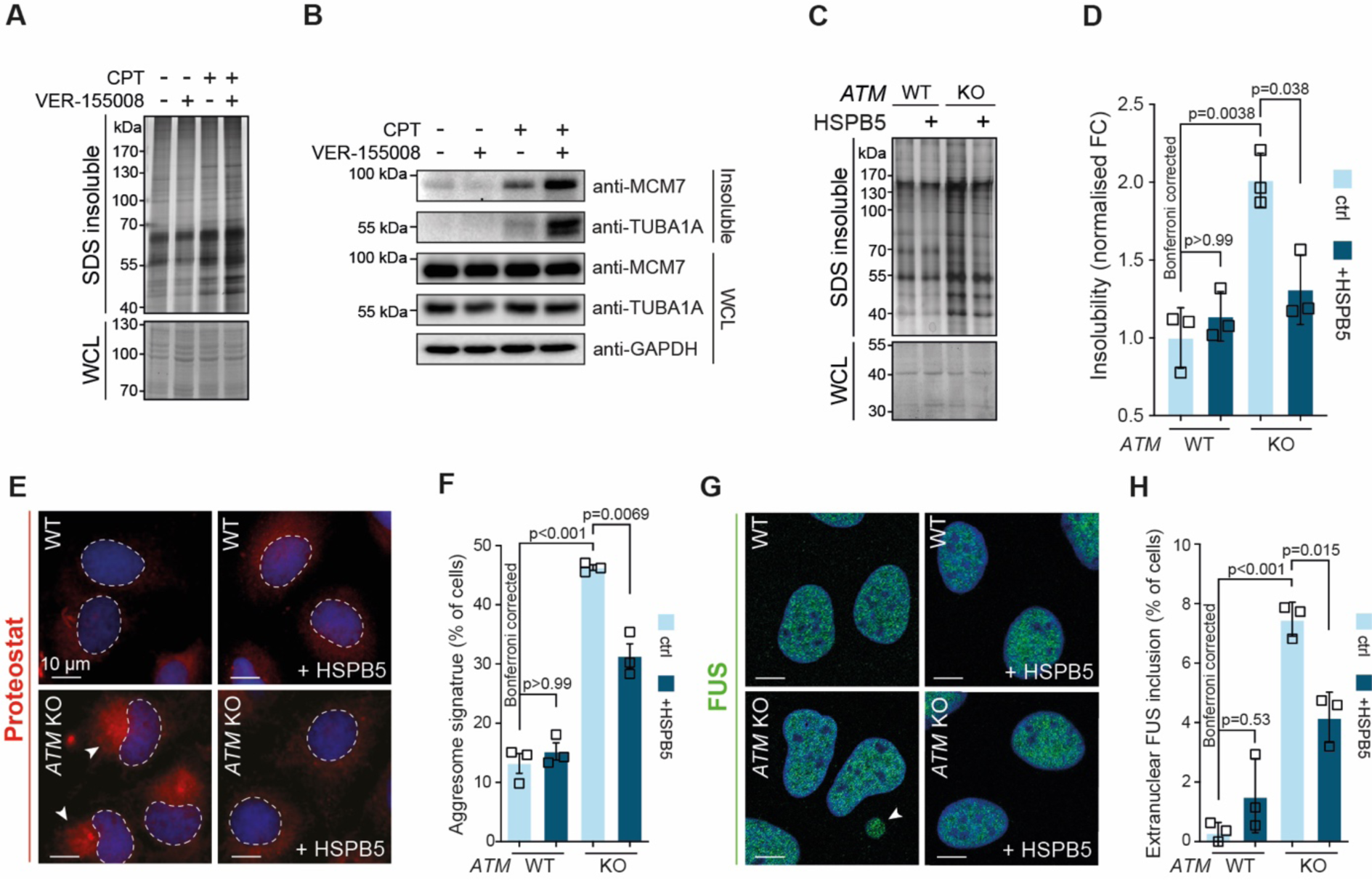
Protein aggregation triggered by genotoxic stress is amenable to modulation by chaperones. See also Figure 6 – figure supplement 1. (A) Aggregated (silver stain) and whole cell lysate (WCL; Coomassie) fractions of HEK293T cells treated transiently with CPT, followed by treatment with the VER-155008 HSP70 inhibitor. n=3. (B) Western blot of WCL and aggregated proteins isolated from HEK293T cells treated transiently with CPT, followed by treatment with the VER-155008 HSP70 inhibitor (10 μM), probed with the indicated antibodies. n=3. (C) Aggregated (silver stain) and whole cell lysate (WCL; Coomassie) fractions of U2OS wild-type and *ATM* KO cells, with or without overexpression of HSPB5. (D) Quantification of C. (E) Representative immunofluorescence pictures of U2OS wild-type and *ATM* KO cells stably overexpressing HSPB5 or not, stained with Proteostat® (red) and Hoechst (blue). (F) Quantification of aggresome signatures in E. (G). Representative immunofluorescence pictures of U2OS wild-type and *ATM* KO cells stably overexpressing HSPB5 or not, stained with anti-FUS (green) and Hoechst (blue). (H) Quantification of extranuclear FUS inclusions in G. In D, F and H, squares represent independent experiments, bars represent mean ± SEM. P-values are obtained by two-tailed Student’s t-tests followed by a Bonferroni correction for multiple comparisons.

We next reasoned that increasing chaperone capacity may also raise the aggregation threshold again. We screened an overexpression library of several major chaperone families, including HSPAs, J-domain proteins (JDPs) and small heat shock proteins (HSPBs) for their ability to reduce the increased protein aggregation triggered by genotoxic conditions, using U2OS *ATM* KO cells as a model (Figure 6 – figure supplement 1A). While most of these did not overtly decrease protein aggregation, overexpression of several JDPs reduced protein aggregation, including the generic anti-amyloidogenic protein DNAJB6b (Aprile et al. 2017; Hageman et al. 2010; Månsson et al. 2014). However, we found that the small heat shock protein HSPB5 (or CRYAB, i.e. *α*B-crystallin) was especially effective. HSPB5 is a potent suppressor of aggregation and amyloid formation (Delbecq and Klevit 2019; Golenhofen and Bartelt-Kirbach 2016; Hatters et al. 2001; Webster et al. 2019).

Interestingly, proteins that aggregate in U2OS *ATM* KO cells are enriched ∼3-fold for reported HSPB5 interactors compared to U2OS background aggregation. Moreover, our RNA sequencing analysis revealed that U2OS cells inherently have a >400-fold higher basal expression of HSPB5 than HEK293T cells (Figure 5 – figure supplement 1D). In U2OS *ATM* KO cells, HSPB5 levels are increased even further: it is one of only three transcriptionally upregulated chaperones (Figure 5F), and it is the only chaperone whose abundance is significantly increased in the U2OS *ATM* KO WCL dataset (Figure 5 – figure supplement 1F).

Based on this, we generated U2OS cells that stably overexpress HSPB5 in both wild-type and *ATM*-deficient backgrounds (Figure 6 – figure supplement 1B), and confirmed that this drastically reduced the enhanced protein aggregation in the latter (Figure 6C,D). HSPB5 overexpression also reduced ProteoStat*®* aggresome staining and the occurrence of cytoplasmic FUS puncta, two other markers of a disrupted protein homeostasis (Figure 6E-H) (Neumann et al. 2006; Shen et al. 2011). Although HSPB5 itself has never been linked to genome maintenance, we evaluated whether HSPB5 may mitigate the increased aggregation following a loss of ATM by altering DNA repair capacity. However, we find no indication for this, as the ionizing irradiation-induced DNA lesion accumulation and subsequent resolution as measured by 53BP1 foci formation was not affected by HSPB5 expression, in neither U2OS wild-type nor *ATM* KO cells (Figure 6 – figure supplement 1C,D).

## DISCUSSION

Here, we report that topoisomerase poisoning and functional impairment of ATM or ATR trigger a widespread aggregation of LLPS-prone and supersaturated proteins. Our data show that the aggregation of these metastable proteins is a consequence of an overload of chaperone systems under these genotoxic conditions. This overload of chaperone systems causes protein homeostasis to shift, lowering the cell-intrinsic threshold of protein aggregation. As a result, vulnerable proteins that are largely kept soluble under normal conditions now succumb more readily to aggregation. The accelerated aggregation of the model substrates polyQ- and luciferase that occurs in cells exposed to these conditions underlines this threshold change.

The observed shift in protein homeostasis after genotoxic stress is strikingly reminiscent of what is believed to occur under conditions of (chronic) stress (Weids et al. 2016), and during many age-related neurodegenerative disorders (David et al. 2010; Hipp et al. 2019; Morley et al. 2002). Supersaturated proteins have been found to be over-represented in cellular pathways associated with these disorders (Ciryam et al. 2015), and disease-associated aggregating proteins, including FUS, tau and *α*-synuclein are known to exhibit LLPS behavior (reviewed in Zbinden et al., 2020). Indeed, we find that the proteins that aggregate in our experiments show a strong overlap in identity and function with stress-induced aggregation, and with the aggregation observed in various proteinopathies.

The shift in protein homeostasis under genotoxic stress conditions can theoretically be caused by either an altered capacity of protein quality control systems, or by an increased demand emanating from an altered proteome. These are however difficult to disentangle fully, in particular because they may form a vicious cycle of events, where (co)chaperones are increasingly sequestered as a growing number of proteins succumbs to aggregation (Klaips et al. 2018). Either way, both result in a net lack of protein quality control capacity, which can be rescued by upregulating specific chaperones, and exacerbated by further decreasing chaperone capacity. The aggregation that occurs under the genotoxic conditions used in our study follows this pattern. Nevertheless, our data point out that at least part of the overload of chaperone systems follows upon an increased proteome demand. Multiple (co)chaperones that have been reported to interact frequently with the aggregating proteins are upregulated under the genotoxic conditions used in our study. Many of these (co)chaperones aggregate themselves as well. Crucially, further overexpression of one of the most upregulated chaperones in U2OS, HSPB5, is able to largely bring aggregation in *ATM* KO cells back down to the wild-type level. The strong overlap between CPT-induced aggregation in HEK293T cells and background aggregation in U2OS cells also argues for an increased demand. U2OS is a cancer cell line (osteosarcoma), whereas HEK293 cells have a vastly different origin (embryonic kidney). Cancer cells inherently exhibit elevated levels of protein stress, which has been attributed to an increased protein folding and degradation demand (Dai, Dai, and Cao 2012; Deshaies 2014). The notion that the rewiring of chaperone systems in response to CPT-treatment in HEK293T cells mimics the difference in chaperone wiring between HEK293T and U2OS cells underlines this further.

Our data indicate that any increased demand caused by these genotoxic conditions is independent of quantitative changes of the aggregating proteins themselves, and likely also of any genetic alterations in their coding regions (*in cis* genetic alterations). The accelerated polyQ aggregation – not accompanied by any enhanced CAG repeat instability – provides support for this. This is not necessarily surprising, as proteins that aggregate as a consequence of an overload of the protein quality control do not have to be altered themselves. Previous studies have shown that during proteomic stress a destabilization of the background proteome can result in a competition for the limited chaperone capacity available, causing proteins that are highly dependent on chaperones for their stability and solubility to aggregate readily (Gidalevitz et al. 2010; Gidalevitz, Prahlad, and Morimoto 2011). In this light, it is interesting that the proteins that aggregate in our experiments are in general prone to engage in liquid-liquid phase separation. Although LLPS is a different biochemical process than protein aggregation (with different underlying mechanisms and principles), aberrant LLPS can drive the nucleation of insoluble (fibrillar) protein aggregates, for example for polyQ (Peskett et al. 2018). It is therefore believed that LLPS events need to be closely regulated and monitored to prevent aberrant progression into a solid-like state (Alberti and Dormann 2019). Although data is so far limited, chaperones, and in particular small heat shock proteins, have been reported to play a pivotal role in the surveillance of biomolecular condensates. For example, the HSPB8-BAG3-HSPA1A complex has been found to be important for maintaining stress granule dynamics (Ganassi et al. 2016), and recent work uncovered that HSPB1 is important to prevent aberrant phase transitions of FUS (Liu et al. 2020). We find that HSPB1, HSPB8, BAG3 and HSPA1A are all upregulated in HEK293T cells treated with CPT. Our data thus point at the possibility that genotoxic stress conditions can exacerbate the normally occurring protein aggregation by increasing the risk of aberrant progression of LLPS processes. The molecular details of this process, and whether or not HPSB5 in U2OS cells acts also at this level remain to be investigated. Interestingly, although HSPB5 itself has so far not been shown to undergo LLPS, like HSPB1, it has been found to associate with nuclear speckles (van den IJssel et al. 1998), which are membraneless as well. HSPB5 has also been shown to be important to maintain the stability of the cytoskeleton (Ghosh, Houck, and Clark 2007; Golenhofen et al. 1999; Yin et al. 2019), and many proteins that aggregate in U2OS *ATM* KO cells are cytoskeleton (-related) components. A growing body of evidence indicates that cytoskeleton organization is regulated through LLPS processes (reviewed in Wiegand and Hyman 2020).

The increased protein aggregation that occurs after a loss of ATM – including in A-T patient brains – has been recently attributed to an accumulation of DNA damage (Lee et al. 2021). As an impaired response to DNA damage is believed to be the primary driving force of A-T phenotypes (Shiloh 2020), these findings have fueled the idea that a disruption of protein homeostasis may be an important disease mechanism in A-T. Our data provide further support for this, as they show that the widespread aggregation caused by a loss of ATM follows a predictable pattern that overlaps strikingly with the aggregation that is believed to underlie many neurodegenerative disorders. Importantly, our findings also provide a proof of principle that other genotoxic conditions – including chemotherapeutic topoisomerase poisons – can have a very similar impact. This points at the existence of a broader link between DNA damage and a loss of protein homeostasis. Although further research is needed to determine the full breadth and relevance of this link, our work may thus offer clues as to why besides impairments in ATM, many other genome maintenance defects are characterized by often overlapping (neuro)degenerative phenotypes as well (Petr et al. 2020).

## MATERIALS AND METHODS

### Statistical analyses

Statistical testing was performed using Graphpad Prism software, except for LFQ proteomics and RNA sequencing, which were analyzed in R (see their respective sections further down for more information). The statistical tests that were used are indicated in each figure legend. For experiments with pairwise comparisons, two-tailed student’s unpaired t-test was used, unless otherwise indicated. For experiments with multiple comparisons, a Kruskall-Wallis with Dunn’s post-hoc test (when datasets did not pass normality testing), or two-tailed student’s unpaired t-tests with Bonferroni correction (when indicated) was performed. P-values are shown for all experiments. All repetitions (n) originate from independent replicates.

### Mammalian cell culture

All cell lines were cultured in DMEM (GIBCO) supplemented with 10% FBS (Sigma Aldrich), 100 units/ml penicillin and 100 µg/ml streptomycin (Invitrogen). HEK293 cells expressing inducible GFP-Htt^exon1-Q71^ (GFP-Q71) have been described previously (Kakkar et al. 2016), and HEK293 cells expressing inducible luciferase-GFP as well (Hageman et al. 2011). U2OS and HEK293 *ATM* KO cells were generated using the ATM CRISPR/Cas9 KO and ATM HDR plasmids (sc-400192, sc-400192-HDR from Santa Cruz) according to the manufacturer’s guidelines. Individual clones were picked and verified by PCR and Western blotting.

### Western blotting and (immuno)staining

For Western blotting, proteins were transferred to either nitrocellulose or PVDF membranes, probed with the indicated antibodies, and imaged in a Bio-Rad ChemiDoc imaging system. For an overview of all antibodies used in this study, see Table 1.

For (immuno)staining, cells were grown on coverslips, fixed in 2% formaldehyde, permeabilized with 0.1% Triton-X100 and incubated for 15 minutes with 0.5% BSA and 0.1% glycine solution in PBS. ProteoStat® staining (ENZO, ENZ-51023-KP050) was performed according to the manufacturer’s instructions. Primary antibody incubation (see Table 2) was performed overnight at 4°C. After secondary antibody incubation cells were stained with Hoechst (Invitrogen, H1399) and mounted on microscopy slides in Citifluor (Agar Scientific). Cells were observed using a confocal scanning microscope (Leica), and images were analyzed using ImageJ software. The aggresome signature was defined as cells exhibiting both a curved nucleus and perinuclear presence of ProteoStat® dye. For DNA repair kinetics experiments, cells were irradiated with 2 Gy (IBL-637 irradiator, CIS Biointernational), fixed at the indicated timepoints post-irradiation, and stained for 53BP1. Images were analyzed using ImageJ software.

**Table 2:**
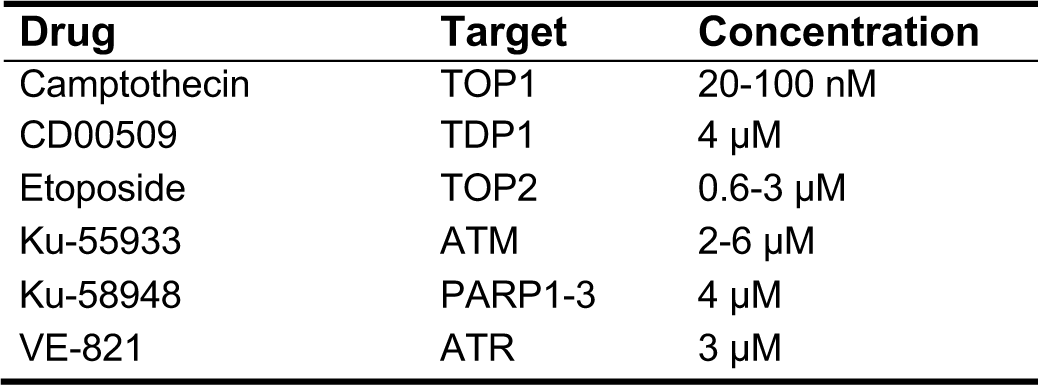
Genotoxic drugs used in this study

### Genotoxic drug treatments

For genotoxic drug treatments, experiments cells were treated with drugs in the indicated doses. The culture medium was replaced 24 hours after drug treatment, and after another 48 hours cells were harvested by scraping in PBS, centrifugation and snap-freezing in liquid nitrogen. See Table 2 for an overview of the drugs and concentrations used in this study.

### Differential detergent protein fractionation

Cells were resuspended in ice-cold lysis buffer containing 25 mM HEPES pH 7.4, 100 mM NaCl, 1 mM MgCL_2_, 1% v/v Igepal CA-630 (#N3500, US Biological), complete EDTA-free protease inhibitor cocktail (Roche Diagnostics), and 0.1 unit/µl Benzonase endonuclease (Merck Millipore) and left for 1 hour on ice with intermittent vortexing. Protein content was measured and equalized, and Igepal CA-630 insoluble proteins were pelleted by high-speed centrifugation (21,000 *rcf*, 45 minutes, 4 °C). Protein pellets were washed with lysis buffer without Igepal CA-630, and re-dissolved in lysis buffer supplemented with 1% v/v SDS at RT in a Thermomixer R (Eppendorf) at 1200 *rpm* for 1-2 hours. SDS insoluble proteins were then pelleted by high-speed centrifugation (21,000 *rcf*, 45 minutes). SDS insoluble protein pellets were washed with lysis buffer without any detergent. For subsequent silver staining, pellets were solubilized in urea buffer (8 M urea, 2% v/v SDS, 50 mM DTT, 50 mM Tris/HCl pH 7.4) overnight at RT in a Thermomixer R (Eppendorf) at 1200 *rpm*. For subsequent Western blotting, pellets were solubilized in sample buffer (), boiled for 10 minutes, and left overnight RT in a Thermomixer R (Eppendorf) at 1200 *rpm*. Fractions were separated using SDS-PAGE, imaged using a Bio-Rad ChemiDoc imaging system, and analyzed using ImageJ software.

### LC-MS/MS Analysis

Samples were reduced (Dithiothreitol 25 mM, 37 °C, 30 minutes), alkylated (Iodoacetamide 100 mM, room temperature, 30 minutes in darkness) and trypsin digested on S-trap columns (Protifi) using the high recovery protocol (http://www.protifi.com/wp-content/uploads/2018/08/S-Trap-micro-high-recovery-quick-card.pdf). After elution, samples where dried up on speed-vac and resuspended in 25 uL of 0.1 % (v/v) formic acid in water (MS quality, Thermo). Mass spectral analysis was conducted on a Thermo Scientific Orbitrap Exploris. The mobile phase consisted of 0.1 % (v/v) formic acid in water (A) and 0.1 % (v/v) formic acid in acetonitrile (B). Samples were loaded using a Dionex Ultimate 3000 HPLC system onto a 75 um x 50 cm Acclaim PepMapTM RSLC nanoViper column filled with 2 µm C18 particles (Thermo Scientific) using a 120-minute LC-MS method at a flow rate of 0.3 µL/min as follows: 3 % B over 3 minutes; 3 to 45 % B over 87 minutes; 45 to 80 % B over 1 minute; then wash at 80 % B over 14 minutes, 80 to 3 % B over 1 minutes and then the column was equilibrated with 3 % B for 14 minutes. For precursor peptides and fragmentation detection on the mass spectrometer, MS1 survey scans (m/z 200 to 2000) were performed at a resolution of 120,000 with a 300 % normalized AGC target. Peptide precursors from charge states 2-6 were sampled for MS2 using DDA. For MS2 scan properties, HCD was used and the fragments were analyzed in the orbitrap with a collisional energy of 30 %, resolution of 15000, Standard AGC target, and a maximum injection time of 50 ms.

MaxQuant version 1.6.7.0 was used for peptides and protein identification (Tyanova, Temu, and Cox 2016) and quantification with a proteomic database of reviewed proteins sequences downloaded from Uniprot (08/17/2020, proteome:up000005640; reviewed:yes). Abbreviated MaxQuant settings: LFQ with minimum peptide counts (razor + unique) ≥ 2 and at least 1 unique peptide; variable modifications were Oxidation (M), Acetyl (Protein N-term), and Phospho (STY); Carbamidomethyl (C) was set as a fixed modification with Trypsin/P as the enzyme.

ProteinGroup.txt from MaxQuant output was used for protein significance analysis via post-processing in R: potential contaminant and reversed protein sequences were filtered out, partial or complete missing values in either case or control replicates were imputed (Dou et al. 2020) using a single seed, log_2_ transformed LFQ intensities were used for t-tests, including Benjamini-Hochberg corrected, p-adjusted values. Log_2_ fold-change for each protein record was calculated by subtracting the average log_2_ LFQ intensity across all replicates in control samples from the average log_2_ LFQ intensity across all replicates in case samples. To mitigate imputation-induced artifacts among significant proteins, only significant proteins detected and quantified in at least two replicates were considered: p-adjusted value ≤ 0.05 and, for cases (log_2_ fold-change ≥ 1, replicates with non-imputed data ≥ 2), or for controls (log_2_ fold-change ≤ −1, replicates with non-imputed data ≥ 2).

### RNAseq library construction and sequencing

RNA was isolated from cells with the AllPrep DNA/RNA Mini Kit from Qiagen. RNA concentrations were measured on a Nanodrop. 150 ng of RNA was used for library preparation with the Lexogen QuantSeq 3’ mRNA-Seq Library Prep Kit (FWD) from Illumina. Quality control of the sequencing libraries was performed with both Qubit™ (DNA HS Assay kit) and Agilent 2200 TapeStation systems (D5000 ScreenTape). All libraries were pooled equimolar and sequenced on a NextSeq 500 at the sequencing facility in the University Medical Center Groningen, Groningen, the Netherlands.

Data preprocessing was performed with the Lexogen Quantseq 2.3.1 FWD UMI pipeline on the BlueBee Genomics Platform (1.10.18). Count files were loaded into R and analyzed with edgeR ^107^. Only genes with > 1 counts in at least 2 samples were included in the analysis. Count data was normalized using logCPM for Principal Component Analysis (PCA). Differential gene expression analysis was performed using the likelihood ratio test implemented in edgeR. Cutoffs of an absolute log fold change > 1 and an FDR-adjusted p-value < 0.05 were used to identify significantly differentially expressed genes.

### Quantification of polyglutamine aggregation

24 hours after seeding, stable tetracycline-inducible HTT Q71-GFP-expressing HEK293 cells were treated with the indicated genotoxic drugs listed in Table 1, as described. Cell lysis, polyQ filter-trap and immunodetection were performed as described previously (Kakkar et al. 2016), and results were analyzed using ImageJ software.

### CAG repeat length analysis

DNA was isolated from HTT Q71-GFP-expressing HEK293 cells through MasterPure™ Complete DNA and RNA Purification Kit (Epicentre®) according to the manufacturer’s instructions. The CAG repeat length analysis was performed by PCR with 100ng of DNA in a 10 µl reaction volume containing AmpliTaq Gold® Fast PCR Master Mix (Applied Biosystems), and 0.2 µM of both forward (HEK293TQ71F [FAM]: 5’ - GAGTCCCTCAAGTCCTTCC - 3’) and reverse (HEK293TQ71R: 5’ - AAACGGGCCCTCTAGACTC - 3’) primers, flanking the CAG repeat tract. The samples were subjected to an initial denaturation step (95° C, 10 min), 35 amplification cycles (96° C, 15 s; 59.2° C, 15 s; 68° C, 30 s) and a final extension of 72° C, 5 min. PCR was followed by capillary electrophoresis in a ABI3730XL Genetic Analyzer, and results were analyzed through GeneMapper Software V5.0 (both Applied Biosystems).

### Retroviral overexpression of HSPB5

Retrovirus was produced in the Phoenix-AMPHO retroviral packaging cell line using a pQCXIN–HSPB5 vector as described before (Vos et al. 2010). U2OS wild-type and *ATM* KO cells were infected in the presence of 5 µg/ml polybrene (Santa Cruz). Cells in which the HSPB5 vector integrated successfully were selected using G418, and HSPB5 overexpression was confirmed via Western blotting.

### Online tools and databases used

For an overview of all tools and databases used in this study, see Table 3.

**Table 3:**
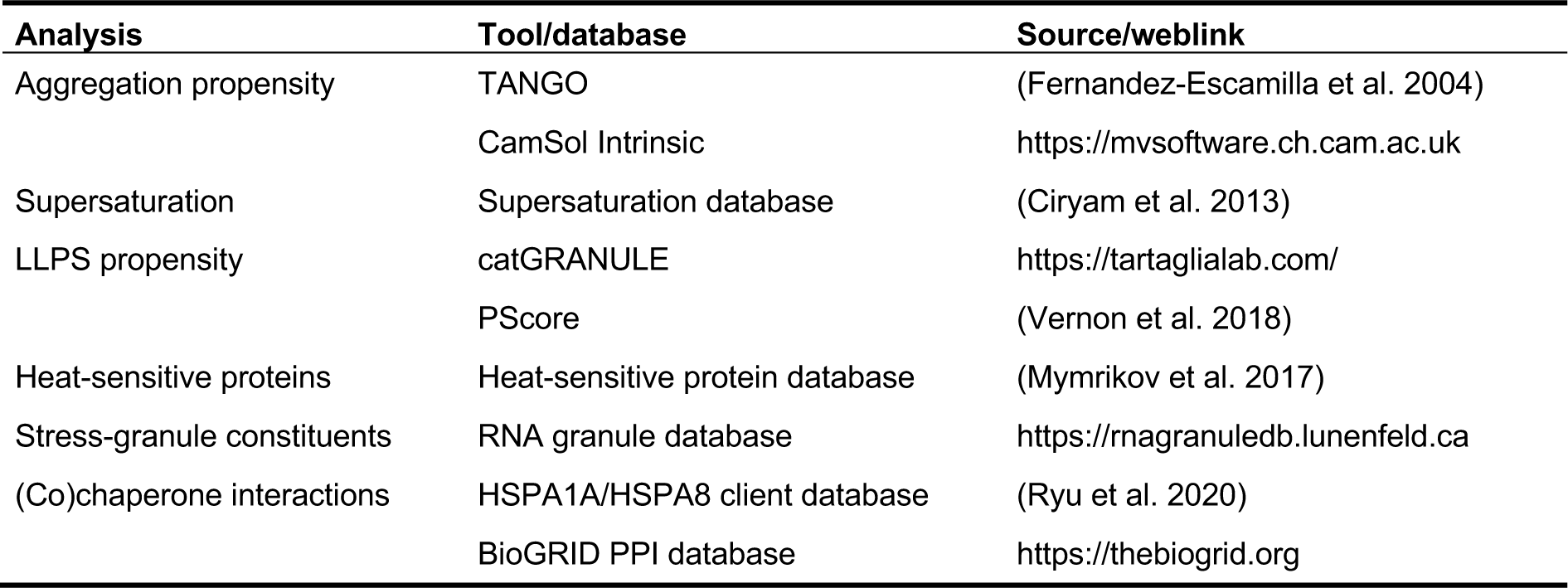
Online tools and databases used

### Data availability

The MS proteomics data have been deposited to the ProteomeXchange Consortium via the PRIDE partner repository (Perez-Riverol et al. 2019) with the dataset identifier PXD025797. The RNAseq data generated in this study are available through Gene Expression Omnibus at https://www.ncbi.nlm.nih.gov.geo with accession number GSE173940. The R codes for MS analysis and for the RNAseq differential expression analysis will be made publicly available through GitHub in the event of acceptance.

## ACKNOWLEDGEMENTS

This work was supported by a Nederlandse organisatie voor Wetenschappelijk Onderzoek (NWO) grant to SB [ALW 824.15.004], and by generous funding from Charity4Brains. This work was partly funded by the National Institute of General Medical Sciences of the National Institutes of Health (NIH) [R01GM126170 to J.L.]; the content is solely the responsibility of the authors and does not necessarily represent the official views of the NIH.

## AUTHOR CONTRIBUTIONS

W.H. and S.B. conceived and designed the study. J.L. and H.H.K. contributed to scientific hypotheses generation. W.H. and J.C.J.v.d.L. performed protein fractionations and Western blotting experiments. L.H.d.S. performed LC/MS-MS work. M.O. performed the MaxQuant analysis of LC/MS-MS data. E.G. performed RNA sequencing and analysis. W.H. and M.K.M. performed all downstream computational analyses. R.M., G.V.F. and M.A.W.H.v.W-V. performed and analyzed filter trap assays, R.M. and G.V.F. performed CAG repeat length analysis. J.C.J.v.d.L generated the luciferase-GFP cell-line, and performed the chaperone screen. S.C. performed the HSP70 inhibition experiments. R.M. generated the HSPB5 overexpressing U2OS cell-lines. J.C.J.v.d.L performed the 53bp1 experiments. W.H. and S.B. interpreted the data. W.H. generated figures. W.H. and S.B. wrote the manuscript. J.L., J.C.J.v.d.L, S.L.D., E.G., L.B. and H.H.K. critically reviewed the manuscript.

## CORRESPONDENCE

Main correspondence to Steven Bergink

Technical proteomics correspondence to John LaCava

## ADDITIONAL FILES

**Figure 1 – figure supplement 1.**
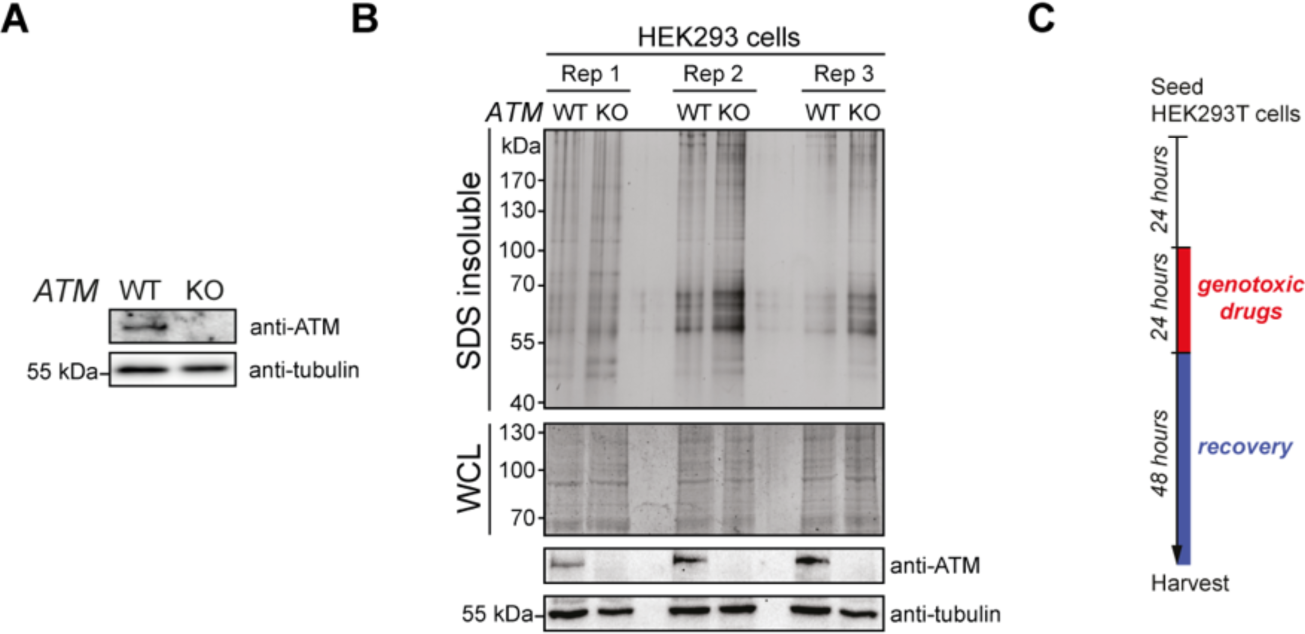
Aggregation is increased in cells lacking ATM. (A) Western blot of U2OS wild-type and *ATM* KO cells, probed using the indicated antibodies. (B) Aggregated (silver stain) and whole cell lysate (WCL; Coomassie) fractions of HEK293 wild-type and *ATM* KO cells. Three independent repeats were loaded on one gel, and stained in-gel. Fractions were also subjected to Western blotting and probed using the indicated antibodies. (C) Experimental outline of Figure 1C and D.

**Figure 2 – figure supplement 1.**
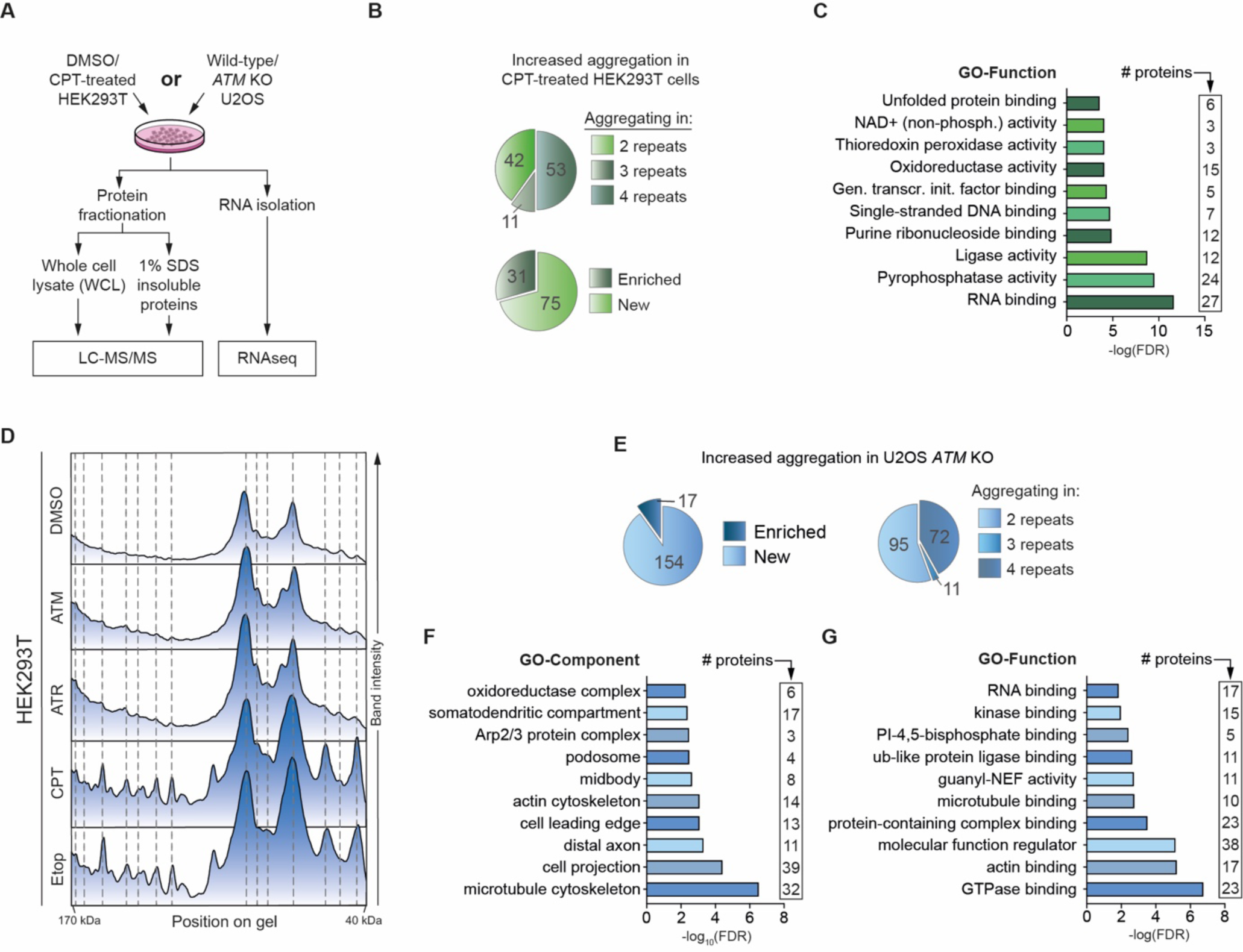
GO-term and densitometry analyses of the aggregation triggered by camptothecin and ATM loss. (A) Experimental outline. All samples were generated in parallel from the same cells, in 4 independent repeats. (B) Top pie chart shows how consistent aggregating proteins were identified across independent repeats. Bottom pie chart shows the relative presence of enriched proteins (i.e. proteins that aggregate more in HEK293T cells treated with CPT) and new proteins (i.e. proteins that aggregate only in HEK293T cells treated with CPT). (C) GO-term analysis (Function) of the increased aggregation in CPT-treated HEK293T cells. The top 10 terms with <2000 background genes are shown. (D) Densitometry analysis of the stained aggregating proteins in Figure 1C. (E) Left pie chart shows the relative presence of enriched proteins (i.e. proteins that aggregate more in U2OS *ATM* KO cells compared to U2OS wild-type cells) and new proteins (i.e. proteins that were only identified as aggregating in U2OS *ATM* KO cells). Right pie chart shows how consistent aggregating proteins were identified across independent repeats. (F,G) GO-term analysis (Component, Function) of the increased aggregation in U2OS *ATM* KO cells. The top 10 terms with <2000 background genes are shown.

**Figure 3 – figure supplement 1.**
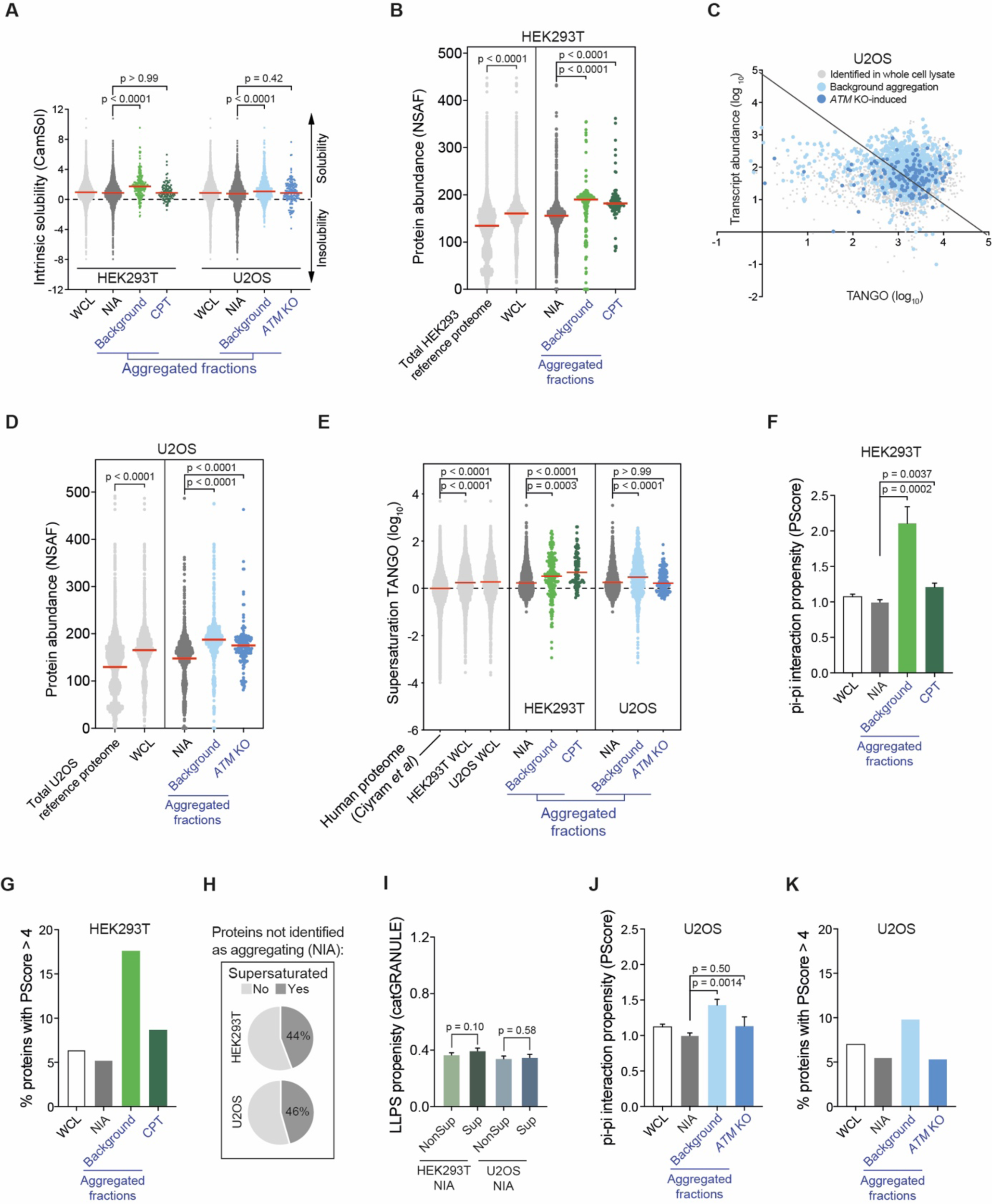
Proteins that aggregate after camptothecin treatment and ATM loss represent a vulnerable subfraction of the proteome. (A) CamSol intrinsic (in)solubility scores of complete WCL, non-aggregated proteins (NIA) and aggregated fractions. HEK293T background and CPT-induced aggregation. Dotted line indicates the theoretical threshold of relative intrinsic (in)solubility. See text for reference. (B) Protein abundances of HEK293T WCL and aggregated fractions obtained by cross-referencing with a HEK293 NSAF (normalized spectral abundance factor) reference proteome, as well as the protein abundances of the entire reference HEK293 reference proteome itself (see text for reference). (C) Transcript abundances (as measured by RNAseq) plotted against TANGO scores, for the complete U2OS MS analysis. All proteins above the diagonal (= U2OS median saturation score, calculated using the U2OS WCL dataset) are relatively supersaturated. See also Figure 3C,D. (D) Protein abundances of U2OS WCL and aggregated fractions obtained by cross-referencing with a U2OS NSAF (normalized spectral abundance factor) reference proteome, as well as the protein abundances of the entire reference U2OS reference proteome itself (see text for reference). (E) Supersaturation scores obtained by cross-referencing with the supersaturation database generated by Ciryam *et al*. (see text for reference). (F) PScores of complete WCL, non-aggregated proteins (NIA) and aggregated fractions in HEK293T. (G) Presence of proteins with a high PScore (>4) in the indicated fractions in HEK293T. (H) Distribution of supersaturated (Sup) and non-supersaturated (NonSup) proteins in the HEK293T and U2OS NIA fractions. (I). CatGRANULE scores of Sup and NonSup proteins in HEK293T and U2OS NIA fractions. (J) PScores of complete WCL, non-aggregated proteins (NIA) and aggregated fractions in U2OS. (I) Presence of proteins with a high PScore (>4) in the indicated fractions in U2OS. For all graphs, circles represent individual proteins, bars represent mean ± SEM. P-values are obtained by Kruskall-Wallis tests followed by Dunn’s correction for multiple comparisons, except in I, where two-tailed Mann-Whitney tests were used.

**Figure 4 – figure supplement 1.**
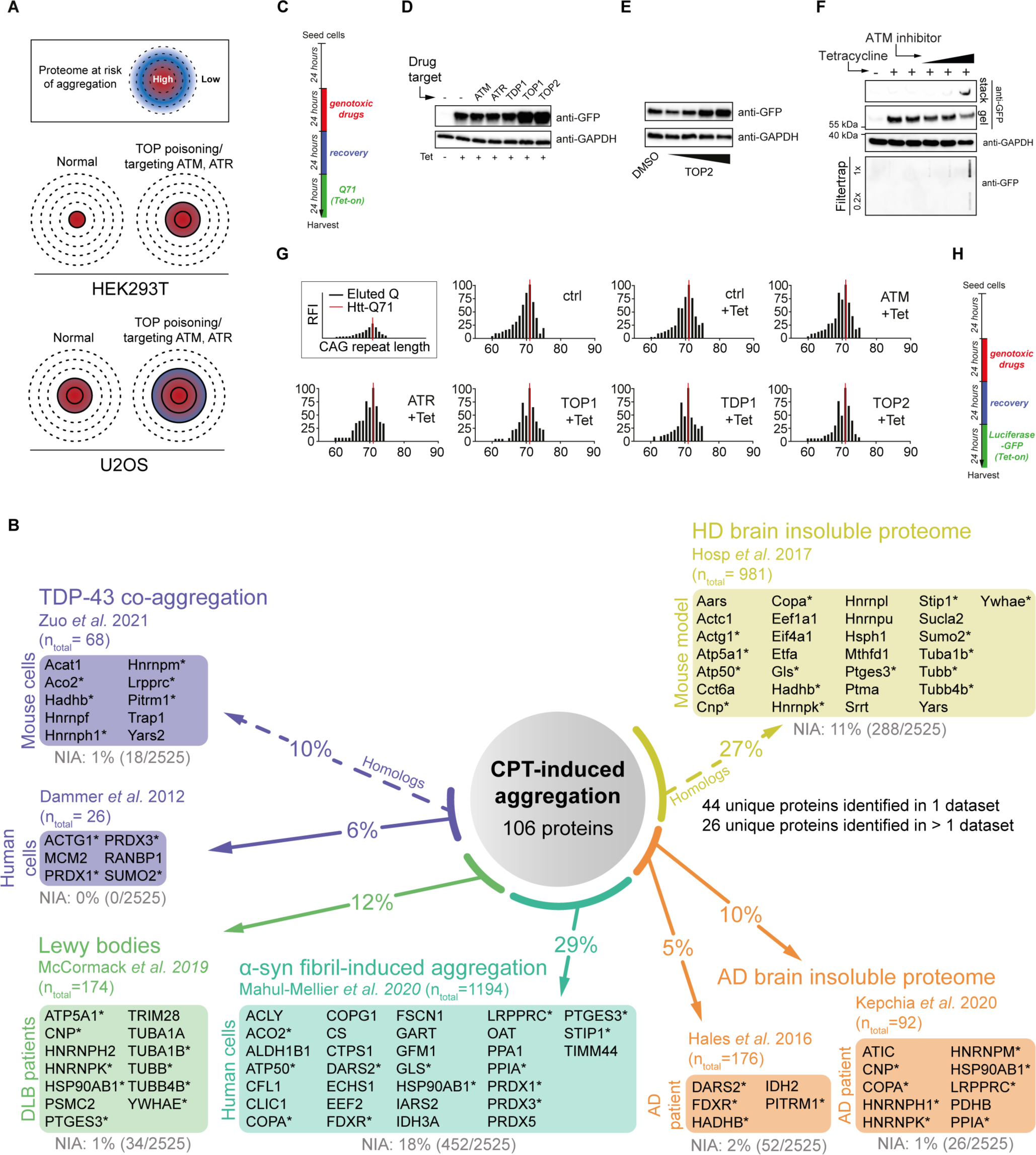
Increased aggregation triggered by camptothecin treatment and ATM loss overlaps with that occurring in various proteinopathies. (A) Conceptual overview of aggregation in HEK293T and U2OS cells. U2OS cells have an inherently lower aggregation threshold, causing more proteins to be affected by aggregation already in the background. (B) Relative occurrences of NIA and aggregating proteins or their mouse homologs in various proteinopathy (model) datasets, obtained from the indicated studies. See also Figure 4D. (C) Experimental outline of Figure 4E and F, and of Figure 4 – figure supplement 1D-F. (D,E) Western blot loading controls of Figure 4E and F, using the indicated antibodies. (F) Western blot and filter trap assay of HEK293 cells expressing inducible Q71-GFP, treated with incremental doses of ATM inhibitor, using the indicated antibodies. n=2. (G). Histograms showing the distribution of CAG repeat length of HEK293 GFP-Q71 cells treated as in C. n=3. (H) Experimental outline of Figure 4G.

**Figure 5 – figure supplement 1.**
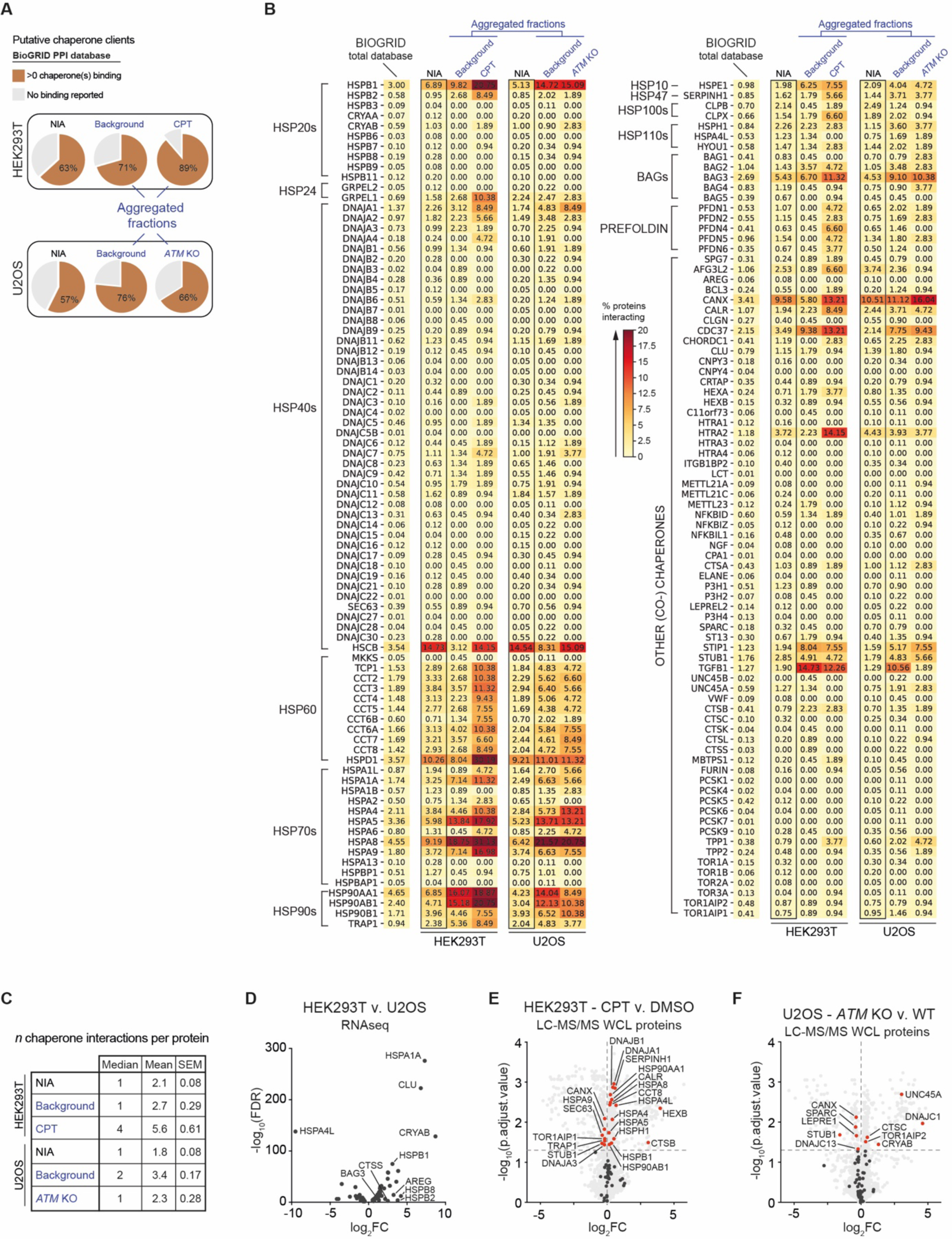
Chaperone systems are rewired in line with the presence of chaperone clients in aggregates induced by camptothecin or ATM loss. (A) Pie charts showing the presence of (co)chaperone interactors (i.e. putative clients) in HEK293T and U2OS aggregated fractions, compared to clients present in both NIA fractions. Interaction data obtained from BioGRID (https://thebiogrid.org). (B) Complete overview of BioGRID (co)chaperone interactions with the aggregated proteins identified in this study, per (co)chaperone. Darker colors represent a higher percentage of proteins with a reported binding to that (co)chaperone. (C) Table showing the number of (co)chaperones logged in BioGRID as interacting with NIA and aggregating protein fractions. (D) Differentially expressed (co)chaperones in U2OS cells compared to HEK293T cells, based on RNA sequencing data. (E,F) Differential expression of (co)chaperones as identified via MS analysis, for HEK293T CPT v. DMSO treated, and U2OS *ATM* KO v. wild-type, respectively.

**Figure 6 – figure supplement 1.**
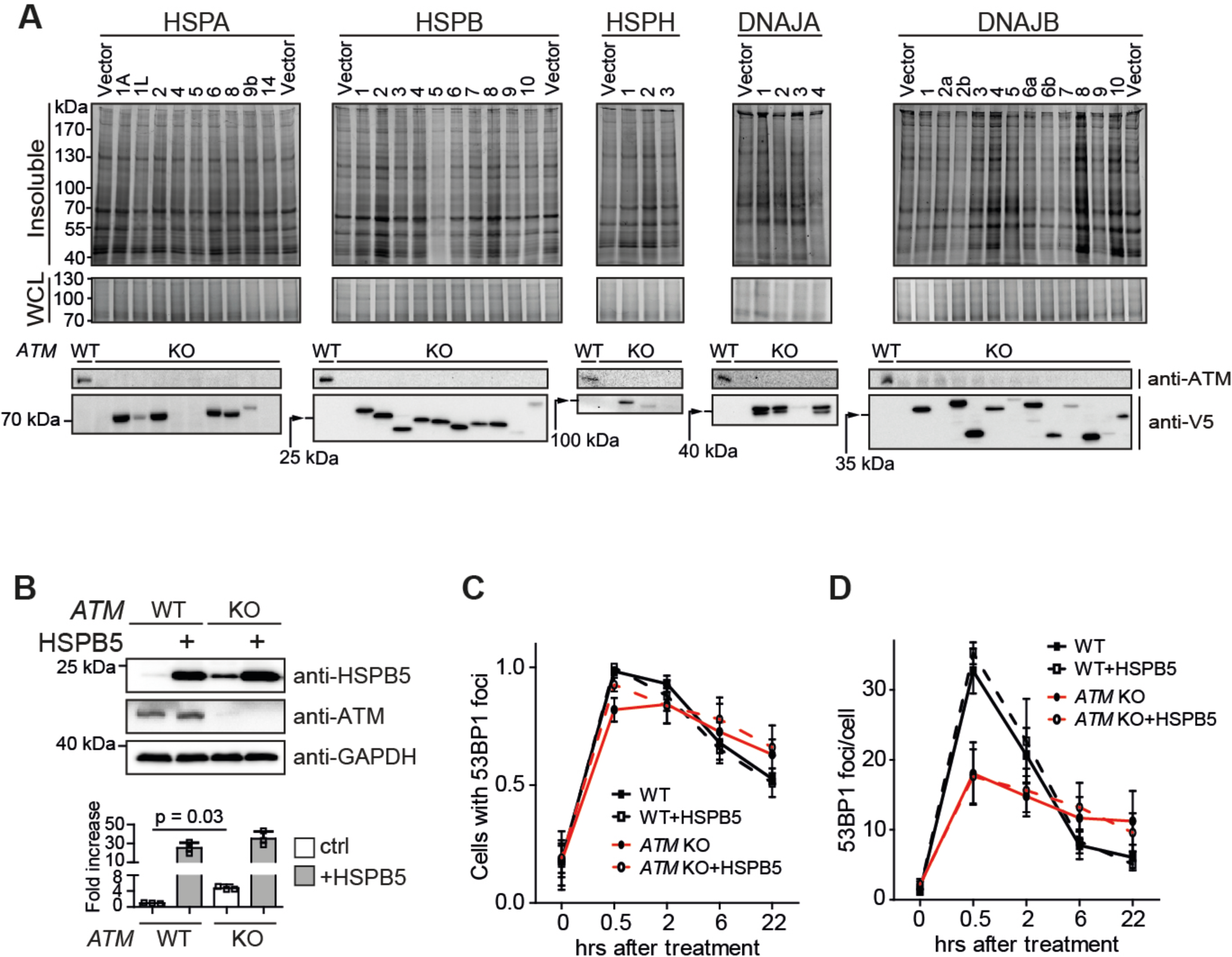
HSPB5 alleviates protein aggregation triggered by a loss of ATM in U2OS cells independent of overt DNA repair capacity changes. (A) Screen for chaperones that can alleviate the increase in aggregated proteins in U2OS *ATM* KO cells. Indicated V5-tagged chaperones were expressed for 48 h. Cells were fractionated and analyzed. SDS-insoluble and WCL fractions were separated by SDS-PAGE and stained by Coomassie. Underneath a Western blot analysis confirming the overexpression of the indicated chaperones, using the indicated antibodies. Note that not all chaperones were equally well overexpressed. (B) Upper panel: Western blot analysis of U2OS wild-type and *ATM* KO cells stably overexpressing HSPB5 (cell lines generated using retroviral infection, see Methods) or not, probed using the indicated antibodies. Lower panel: Quantification of the upper panel. Mean ± SEM, squares indicate independent experiments. Two-tailed t-test. n=3. (C) Plot showing the number of U2OS wild-type and *ATM* KO cells stably overexpressing HSPB5 or not with 53BP1 foci, after 2 Gy of γ-irradiation. (D) Plot showing the number of 53BP1 foci per cell in C. In C and D, bars represent mean ± SD of at least three independent experiments.

**Supplementary File 1** – Supplemental Table 1: MS datasets of aggregated and WCL protein fractions

**Supplementary File 2** – Supplemental Table 2: RNA-sequencing differential expression analysis

**Supplemental Table 2. Differential expression analysis of RNA sequencing data**

**ABOUT THIS DOCUMENT**

This document lists all significant differentially expressed (DE) genes that were identified in our RNA sequencing analysis. DE genes were identified using the likelihood ratio test implemented in edgeR.

These data are from 4 independent biological repeats.

Sheet 1 contains DE genes in U2OS *ATM* KO compared to U2OS WT

Sheet 2 contains DE genes in CPT-treated HEK293T cells compared to vehicle-treated (DMSO) HEK293T cells

Sheet 3 contains DE genes in normal, unstressed U2OS cells compared to unstressed HEK293T cells

